# NMR Hydrogen Exchange and Relaxation Reveal Positions Stabilized by p53 Rescue Mutants N239Y and N235K

**DOI:** 10.1101/2021.02.26.433080

**Authors:** Jenaro Soto, Colleen Moody, Ali Alhoshani, Marilyn Sanchez-Bonilla, Daisy Martinon, Melanie J Cocco

## Abstract

Inactivation of p53 is found in over 50% of all cancers; p53 disfunction is often caused by a single missense mutation localized in the DNA binding domain (DBD). Rescue mutants N235K and N239Y stabilize and restore function to multiple p53 cancer mutants. Here, we use NMR to compare protein dynamics between WT and rescue mutants to understand the mechanism of stabilization. We measured and compared folding dynamics by calculating protection factors (PFs) from NMR hydrogen exchange rates of backbone amides. We find that both rescue mutants impose a global stabilizing effect that dampens their motions compared to WT DBD, predominantly in the β-sandwich. However, a few regions become more flexible in rescue mutants. Notably, positions that have increased PFs map to cancer mutants rescued by each mutant. We also compared relaxation results to obtain flexibility information in the ps to ns timescale regime. Protein sequence analysis was used to determine the occurrence of these rescue mutants in nature and showed that 235K is found in mice and rats, but there is no evidence of 239Y occurring naturally in any species. Understanding the mechanism by which stabilizing mutants rescue p53 may reveal novel avenues for the development of cancer therapeutics. Our findings suggest that cancer therapeutics aimed at restoring p53 function could consider protein dynamics as a metric of drug efficacy.

**Statement of Significance:** Two mutations (N235K and N239Y) within the DNA binding domain of p53 are known to reverse the effects of multiple cancer mutants. However, the mechanism of rescue is not clear since the crystal structures of these mutants are virtually identical to that of the WT. Here we use NMR methods to show that the protein dynamics of the rescue mutants are significantly different compared to WT, allowing us to describe the stability on a per-residue basis. Although these mutations are only four residues apart, they stabilize the structure distinctly in different regions, consistent with the specific cancer mutations they to which they restore function.

## Introduction

p53 is a transcription factor involved in apoptosis, senescence, and DNA repair. When inactivated, cells proliferate unchecked, leading to cancerous tumor growth. Mutations in the *TP53* gene can cause inactivation of the p53 protein through a variety of mechanisms. Many of these missense mutations are disruptive due to the low intrinsic stability of the protein. The prevalence of mutated p53 in cancer has motivated researchers to find drugs that could restore function to these destabilized mutants (1, 2).

Some cancer-causing mutations in p53 can be counteracted by introduction of additional point mutations known as rescue mutations (3). N235K and N239Y are rescue mutations capable of restoring function to multiple prevalent p53 cancer mutants (4, 5). This restorative ability of N235K and N239Y has motivated efforts aimed at understanding the mechanism of rescue (3, 5-9). Both rescue mutations maintain or increase the stability compared to that of wild-type (WT) p53. The stability of N235K is virtually the same as WT with a small increase in ΔΔG reported as 0.3 kcal mol^-1^ (8) and 0.19 kcal mol^- 1^ (9). The stability of N239Y has been reported to increase more substantially compared to WT with ΔΔG of 1.49 kcal mol^-1^ (5), 1.37 kcal mol^-1^ (7), 0.9 kcal mol^-1^ (9). The crystal structures for both rescue mutations show that the structures are virtually identical compared to the WT structure **(Figure 1B)** (9). Since the crystal structures do not reveal a structural mechanism for stabilization, we hypothesize that the mode of rescue may be dynamic in nature.

**Figure 1.**
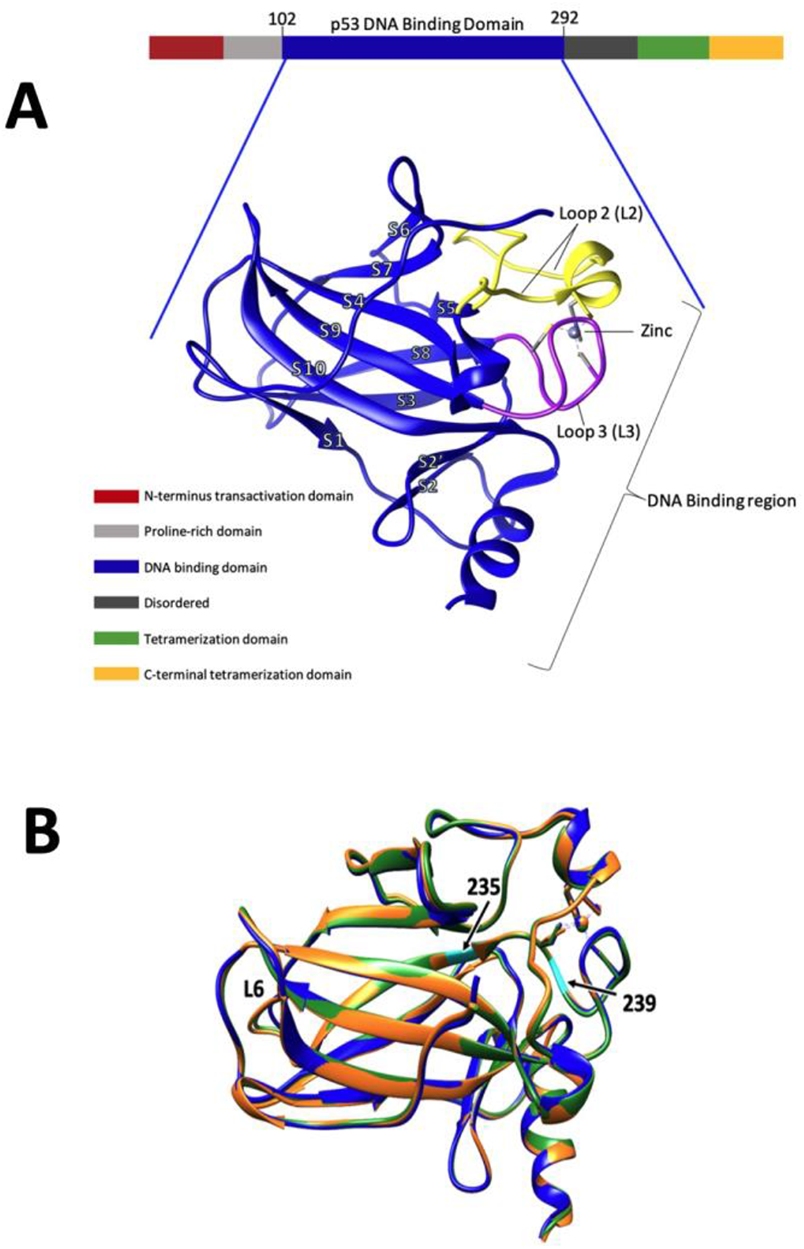
**A)** p53 domains and, highlighted in blue, the structure of the p53 DNA-binding domain (DBD) (PDB: **<u>2XWR)</u>** is shown. Most cancer mutations occur in the DBD. Loop 2 (L2) and loop 3 (L3) are highlighted in yellow and purple respectively [adapted from (11)]. **B)** Overlay of p53 structures: WT (blue, PDB: **<u>2XWR</u>**); N235K (orange, PDB: **<u>4LO9</u>**); N239Y (green, PDB: **<u>4LOE</u>**). Mutation positions 235 and 239 are highlighted in cyan. Only minor changes are seen in side-chain positions near the site of mutation and in L6 in the static structures.

Both cancer and rescue mutants tend to be located within the DNA binding domain (DBD) of p53 (1). Consequently, the majority of dynamic and structural studies have been conducted on the DBD. Furthermore, it has been shown that the DBD plays a major role in the stability of full length p53 (10). The DBD is one of six p53 domains and it is comprised of 190 residues (102-292) **(Figure 1A)**. The secondary structure within p53 includes β-sandwich with Greek key topology, two helices, and a zinc binding region. The β-sandwich consist of two anti-parallel β-sheets with four (S1, S3, S8, and S5) and five (S4, S6, S7, S9, and S10) strands that pack against each other to form a hydrophobic core (11). Understanding the mechanisms that stabilize the DBD structure could guide future therapeutic interventions for cancers related to p53 instability.

Molecular dynamics (MD) simulations (12) and graphical theoretical analysis of DBD crystal structures (13) show p53 cancer mutants to be more flexible than WT. Simulations have also shown that rigidity is restored by rescue mutants (12). Most recently, an MD simulation performed by Pradhan and coworkers shows that cancer mutant conformations undergo opening more readily compared to WT and rescue mutants (14). This ‘open’ conformation is thought to expose a region of the protein that is prone to aggregation thereby promoting deleterious accumulation of p53 aggregates. Together these studies predict that dynamics may play a dominant role in p53 stability and consequently, ability to function.

We used solution state NMR spectroscopy to compare protein dynamics between WT, N235K, and N239Y DBDs. We compared reversible unfolding dynamics by calculating protection factors from backbone amide hydrogen-deuterium exchange rates. In addition, we determined relaxation parameters (R1, R2, NOE) to obtain flexibility information in the ps to ns time scale. Protection factors show that both N235K and N239Y restore function by imposing global stabilizing effect throughout the DBD. Interestingly, the distribution of amino acids stabilized by each rescue mutation were distinct: N235K strongly stabilizes one sheet of the β-sandwich while N239Y results in extensive stabilization of both sheets. Relaxation data showed both rescue mutants effect dynamics of loop region across the DBD, including imposing new flexibility compared to WT. We find that the effects of these suppressor mutations are far-reaching as we see changes that correlate with either more rigid or flexible local structures throughout the DBD.

## Materials and Methods

### NMR data collection and sample preparation

WT and mutant p53 constructs were provided by Rainer Brachman. ^15^N-labeled p53 DBDs were prepared similarly to published methods (15). The DBD core domain of human p53 (94-312) and rescue mutants were transformed into BL21 *E. Coli* cells. Bacteria were grown at 30 °C in ^15^N-enriched Neidharts minimal media to a density of 0.8-1.0 OD_(550 nm)_. The temperature was lowered to 18 °C and both IPTG and ZnCl2 were added to a final concentration of 1 mM. Cells were allowed to grow for an additional 6-8 hours and then harvested by centrifugation and frozen. The cell pellet was resuspended in 20 mM sodium phosphate, pH7.2, 10 mM BME, 0.5 mM PMSF, and lysed using sonication. Cell debris were removed by centrifugation at 4 °C for 30 minutes. The supernatant was applied to a SP-sepharose column and the protein eluted with a gradient of 100 mM - 600mM NaCl. In some cases, additional purification using a Superdex-75 gel filtration column was performed. Samples were dialyzed into 15 mM potassium chloride, 25 mM sodium phosphate, pH 7.1, 10 mM BME and concentrated to a protein concentration of 300-400 μM. 0.5 mM DSS was added as a chemical shift reference.

All NMR experiments were performed on Varian Inova 800 MHz at 20 °C. NMR data were processed using nmrPipe (16). Residues assignments and rate measurements (using peak volumes) in all 2D HSQC spectra were accomplished using CcpNmr Analysis (17).

3D^15^N-TOCSY-HSQC (τ_mix_ = 75 ms) and NOESY-HSQC (τ_mix_ = 100 ms) (18) were analyzed for assignment of backbone amide resonances. Published WT assignments (15) were confirmed in our WT sample and these were compared to spectra of N235K and N239Y. **Table S1** includes mutant assignments that deviate from WT.

Hydrogen-deuterium exchange experiments were performed with 350 μl samples that were lyophilized and fully suspended into an equivalent volume 100% D_2_O immediately before data collection (∼16 min after initiation of exchange). Since our intent is to compare exchange rates for the most stable positions between WT and two mutants, we chose lyophilization as the technique to transfer protein into D2O to ensure that all samples would have the same final conditions (*e*.*g*., [HDO]). Two-dimensional ^1^H-^15^N heteronuclear single quantum coherence (HSQC) NMR spectrum were recorded for each sample every 70 minutes for 24 hours, and then every 180 minutes for 48 hours at 20 °C. HSQC spectra were collected over a period of one month. ^1^H spectra were also collected for S/N calibration using the DSS signal. Sealed samples were stored in a water bath at 20 °C when not in the magnet. Fresh protein samples had an absorbance at 400 nm of 0.05 or less. During the exchange experiment, samples were assessed for aggregation by light scattering and were no longer used if the absorption at 400 nm exceeded 0.1.

*Calculation of chemical shift changes*, Δδ_amide_ was calculated using the changes in chemical shift of ^1^H and ^15^N according to the following equation from Williamson 2013 (19):

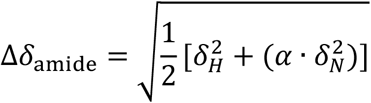

Where *δ*_*H*_ is the change in chemical shift of ^1^H and *δ*_*N*_ is the change in chemical shift of ^15^N between WT and rescue mutant. *α* is the nitrogen scaling factor of 0.14. Residues were considered to have a significant change in chemical shift if they had a Δδ_amide_ > 0.075 (blue line in **Figure 2**). Value 0.075 was calculated by using *δ*_*H*_ *=* 0.05 and *δ*_*N*_ *=* 0.05 consistent with a previous analysis of p53 mutants (15).

**Figure 2.**
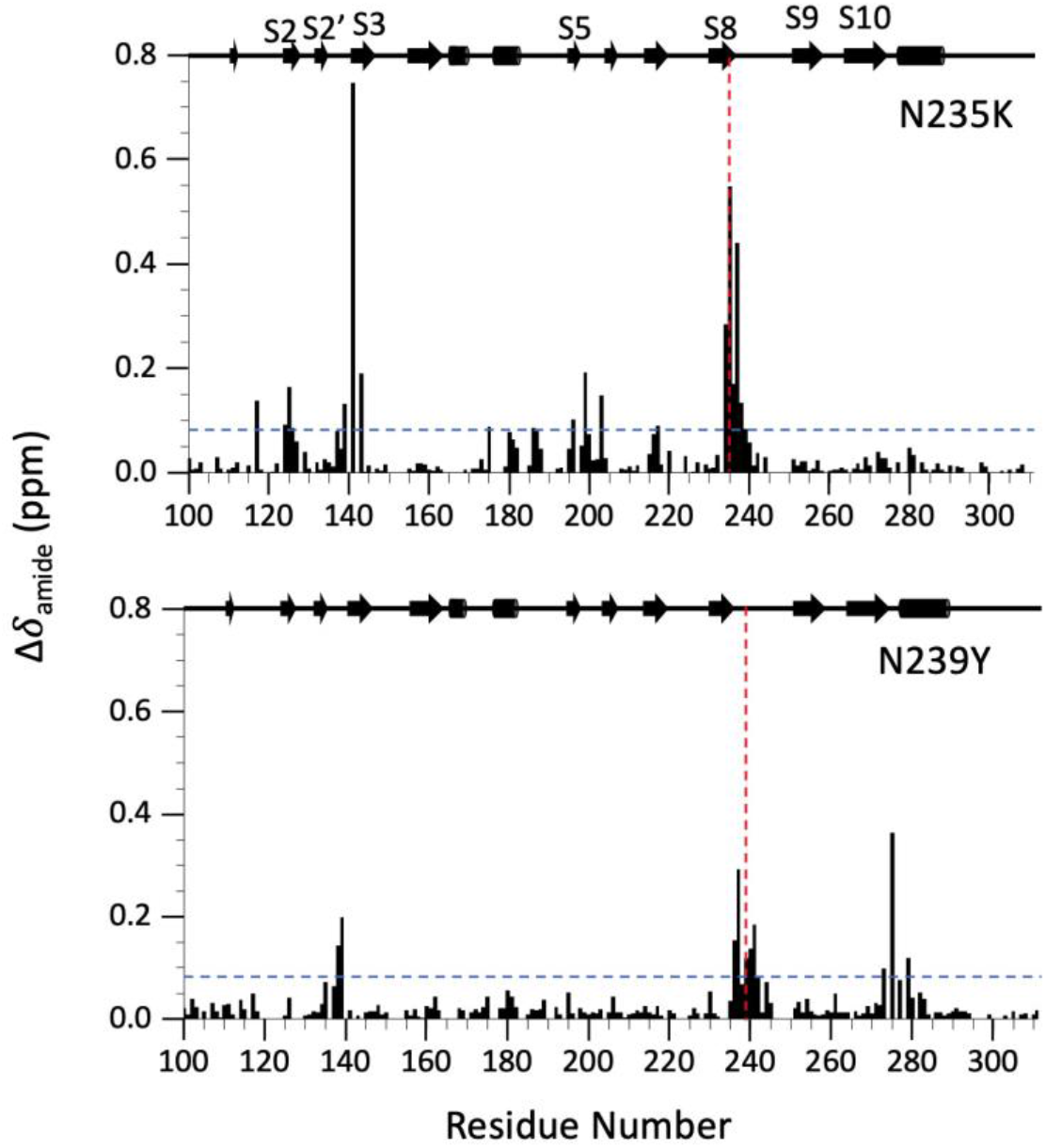
Change in chemical shift Δδamide for residues in N235K (top), and N239Y (bottom) compared to WT. The diagram above illustrates secondary structure elements. Red dashed lines indicate site of mutation. Blue lines indicate changes in chemical shifts that were significant in the evaluation of other p53 mutants (15).

### Relaxation analysis

^15^N longitudinal T1, transverse T2 relaxation and NOEs were measured using pulse sequences of Farrow, et al (20). Spectra were recorded with relaxation delays of 10, 50, 100, 200, 400,750, 1000, 1500 ms for T1 and 10, 30, 50, 70, 90, 110, 130, 150 ms for T2. T1 and T2 times were scrambled during acquisition and 10 ms time spectra were recorded at the beginning and end of both T1 and T2 experiments to confirm S/N had not changed. NOE spectra were recorded with a 5-second irradiation and 3-second delay. R1 and R2 rates were obtain by measuring peak volume as a function of delay time and fitting them to a single exponential function with CcpNmr software (17). Errors in relaxations rates were obtained through the covariance method in CcpNmr.

NMR generalized order parameters (S^2^) were obtained by using *Modelfree 4*.*15* (21, 22) in combination with *FASTModelfree* (23). Initial estimation of *Modelfree* parameters were obtained through the programs *pdbinertia, r2r1_tm, quadric_diffusion* provided on the CoMD/NMR website (24). Residues with NOE and 0.05^2^ values less than 0.6 and 10, respectively, were excluded from *quadric_diffusion* calculation. We found no significant difference for S^2^ results whether *Modelfree analysis* is performed with crystal or NMR structure files (**Figure S1**). Our final analysis used the Protein Data Bank (PDB) file <u>2FEJ</u> for all p53 variants. All *FASTModelfree* analysis used an NH bond length of 1.015 Å and chemical shift entropy (CSA) value of -179 ppm (25) to allow us to compare to published analysis of WT p53 DBD measured at 600 MHz [see figure in 3 (26)]. *FASTModelfree* analysis proceeded until all parameters converged.

### Sequence analysis

The Uniprot database (27) was utilized to select and align p53 sequences from different species. Calculations of sequence identities/similarities were accomplished through the ExPASy alignment program (28).

### Protein structure analysis

UCSF Chimera (29) was used for protein visualization and analysis. Residue distances were calculated by measuring the distance between alpha carbons. Surface area accessibility was calculated with the areaSAS attribute tool from Chimera using amide hydrogens selected from NMR p53 DBD structure 2FEJ.

## Results

### Structural perturbations in solution

Although the crystal structures of WT and rescue mutants are very similar (RMSD = 0.48 Å for N235K/WT and 0.39 Å for N239Y/WT, overlay in **Figure 1B**), NMR chemical shifts may reveal changes to the local environments when the protein is free in solution. To investigate if these rescue mutations change the average solution structure, we performed chemical shift analysis of ^1^H-^15^N HSQC spectra. The spectra for the two p53 variants were well dispersed and our WT spectra were in excellent agreement with published data (15) (**Figure S2**). Peak positions were compared between the published WT assignments and rescue mutant DBDs. We used 3D ^1^H-^15^N TOCSY and NOESY and spectra to confirm assignments in N235K and N239Y and we were able to catalog chemical shifts for 84 % and 82 % of residues, respectively. Several peaks in the mutant spectra experience a significant chemical shift as a result of the residue substitution **(Figure S3, Table S1)**. We calculated the change in chemical shift (Δ*δ*_amide_) of each peak on the ^1^H-^15^N HSQC spectra using equation (8) from Williamson 2013 (19) **(Figure 2)**. Overall, we found that most peaks remained relatively unchanged in both rescue mutants. N235K had an average Δ*δ*_amide_ 0.05 SD of 0.04 0.050.09 ppm, and that of N239Y was 0.03 0.05 0.05 ppm. The peaks with the largest chemical shift changes are located near the site of mutations, including residue 139, 141, 143, 196, 234-238 for N235K, and 139, 273, 275, 279, 235-241 for N239Y. This is expected due to the change in local chemical environment caused by the introduction of the rescue mutation side chain. Lack of chemical shift changes further away from the mutation site suggest that N239Y retains the WT structure similar to what was seen in the crystal structure. In contrast, N235K has two peaks with modest chemical shift changes that are far from the site of mutation. These peaks correspond to residue 117 (Δ*δ*_amide_ =0.14, 19.9 Å) and 125 (Δ*δ*_amide_ = 0.16, 11.5 Å), which are near to each other in the three-dimensional structure of p53 in the vicinity of L1 and S2. Thus, N235K may impose some local structural perturbation in the proximity of this region near the DNA binding interface.

### Backbone exchange of DBD

To compare backbone dynamics that modulate hydrogen-deuterium exchange, we assessed protection factors (PFs) by measuring exchange rates of backbone amides. Amide hydrogens that are readily exposed to solvent will exchange at a higher rate compare to hydrogens that are sequestered away from solvent (*i*.*e*., buried in hydrophobic core or involved in secondary structure). There are two mechanisms that could affect rates of amide exchange: changes in unfolding or a structural change whereby an amide becomes more or less occluded from solvent. Since we know that the crystal structures of WT, N235K and N239Y are virtually superimposable and most chemical shifts are unaffected by these mutations, solvent accessibilities should be very similar. Consequently, differences in the rate of exchange should reflect the influence of the mutation on the local unfolding of that particular amide.

We calculated hydrogen exchange rates (k_ex_) of backbone amides to determine PFs, which gives us a measure of how well the local protein structure protects hydrogen atoms against exchange (30). A stable protein would be expected to have backbone amides with a high PF’s, given that it has a folding/unfolding equilibrium that favors the folded, protected state. We determined k_ex_ by measuring peak volumes as a function of time after transfer into D_2_O and fit the results to calculate the exchange rate **(Figure S4)**. PFs were calculated by k_rc_/k_ex_, where k_rc_ is the exchange rate of a backbone amide in random coil and k_ex_ is the experimental exchange rate calculated from our data. k_rc_ was calculated based on equation 2b in Bai, et. al. (31) **(Tables S2-S4)**.

Obtaining PFs for a wide variety of residues gave us a thorough understandings of DBD dynamics. Our DBD construct is composed of 218 residues (94-312), 18 of which are prolines that do not have a corresponding amide hydrogen. Out the 200 amide signals possible, ∼140 backbone hydrogens exchange with the solvent (D_2_O) within the deadtime for this experiment (∼16 minutes). We were able to obtain hydrogen-deuterium exchange rates for 54 residues of the WT, 48 residues of N235K, and 52 residues of N239Y **(Figure S5, S6, Tables S2-S4)**. Protected residues are found throughout the secondary structures of the entire DBD in all variants (mapped onto the structure in **Figure 3**). We calculated the average PF for the top 10 protected residues for each protein (<PF>_top 10_). N235K had the highest <PF>_top 10_ of 15.9 × 10^6^, follow by N239Y with an <PF>_top 10_ of 9.2 × 10^6^, and lastly the WT with the lowest <PF>_top 10_ of 5.6 × 10^6^. Thirtyfive amides were more protected in N235K than in the WT, and 45 amides had larger protection in N239Y compared to WT. The magnitude of increased PF varied between the two rescue mutants **(Figure S7)**. Each rescue mutant had 18 residues with considerable higher (>2-fold) PF compared to WT. These highly protected amides are found throughout the β-sheet sandwich **(Figure 4)**. N235K has the most protected amide which belongs to 237, with a protection factor of 45.9 ×10^6^. Residue 268 was the highest protected amide for both WT and N239Y, with a protection factor of 17.0 × 10^6^, 23.6 × 10^6^, respectively. In general, we found that the amides in the rescue mutations show a higher PF than the amides in WT, but the distribution is distinct for each rescue mutant.

**Figure 3.**
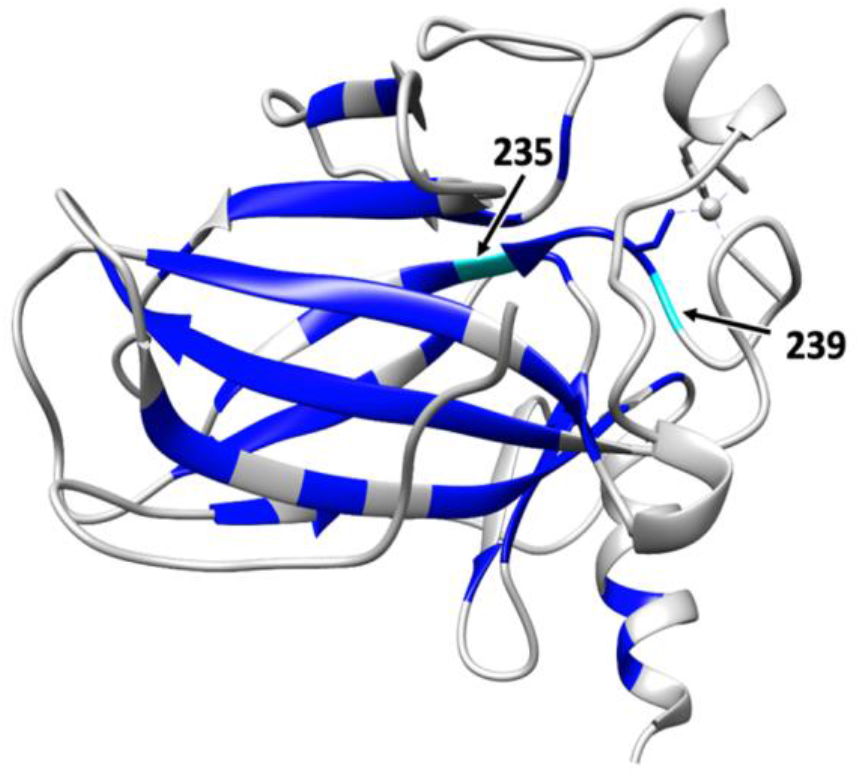
Positions resistant to immediate hydrogen exchange mapped onto the WT p53 structure; protected residues highlighted in blue.

**Figure 4.**
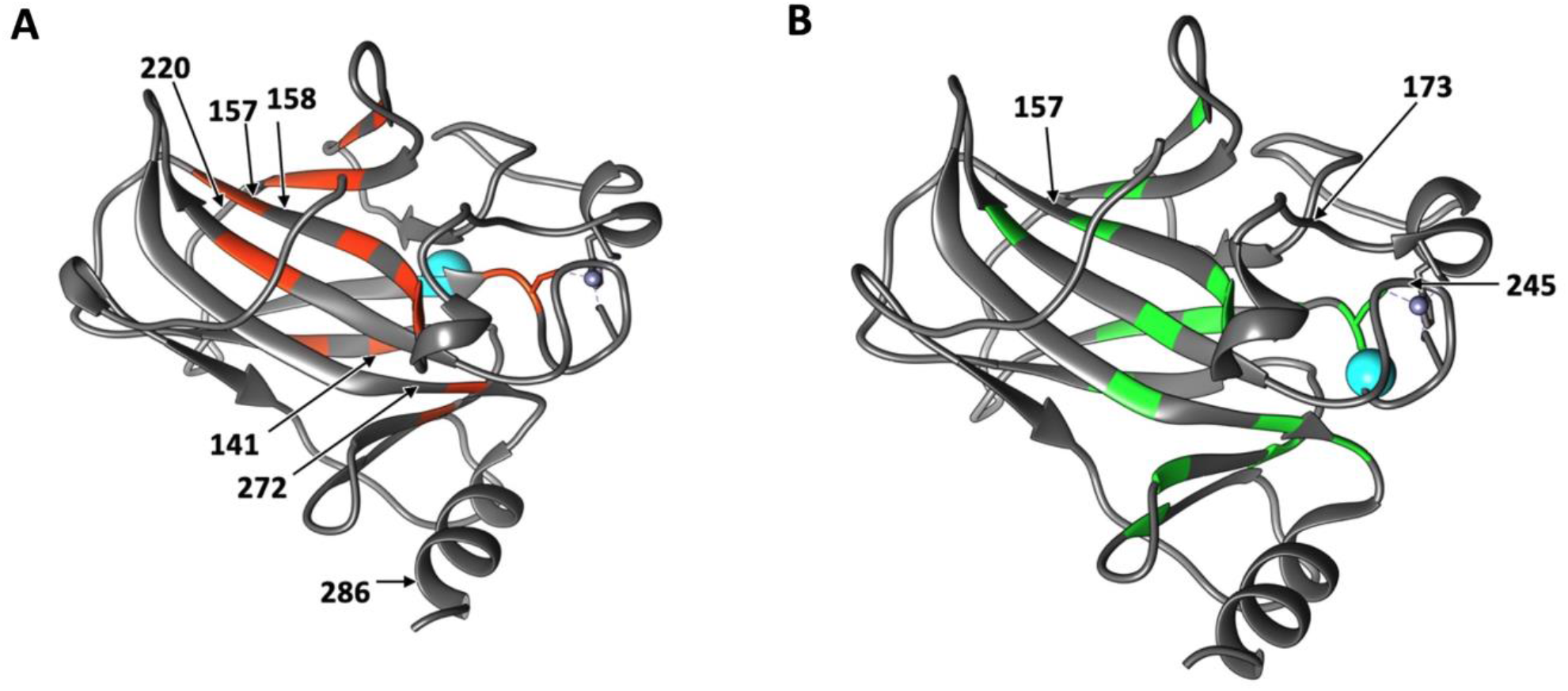
Positions where the rate of amide exchange has slowed at least a two-fold in N235K and N239Y compared to wild-type p53. **A)** 18 backbone hydrogens of N235K more resistant to exchange correspond to residues 134, 141,143, 156, 157, 160, 162, 163, 204, 206, 216, 217, 218, 237, 238, 255, 256, and 273. These residues are highlighted in orange on the structure of the N235K rescue mutation (PDB: **<u>4LO9</u>**). **B)** 18-back-bone hydrogens of N239Y more resistant to exchange correspond to residues 127, 132, 135, 143, 158, 162, 163, 206, 217, 232, 234, 236, 238, 253, 257, 270, 273, and 275. The residues are highlighted in green on the structure of the N239Y rescue mutation (PDB: **<u>4LOE</u>**). Positions of rescue mutations are denoted by cyan spheres. Arrows point out cancer mutation positions known to be rescue by N235K and N239Y, respectively. The pattern of positions with increased protection are different between the two rescue mutants. Notably these mutants rescue different cancer mutations.

PF ratios reveal regions of increased stability in both rescue mutants compared to WT. Although PFs are usually plotted as a function of sequence number, we found it useful to plot the ratio of PFs (mutant/WT) by secondary structure unit. **Figure 5** shows a 3D bar graph organized by β-strand for each β-sheet of the sandwich. We observe that N235K stability is increased on the side of the β-sandwich opposite to the mutation but only two positions on the side containing the mutation. This includes increase protection in strands S4, S6, S7 and parts of S9(254-256) and S10(265-268). We note that N235K also showed regions of decreased protection compared to WT including S3(144-146) and S8(232-236). The same analysis was performed on N239Y; in contrast to N235K, we found that stability is increased on both sides of the β-sandwich in N239Y. All strands have residues with increased protection. In general, we observe that the rescue mutations impose long-range stability throughout the DBD; however, the patterns are distinct: N235K shows a clear trend of asymmetric stability while N239Y stability is more widely distributed throughout both sheets of the β-sandwich.

**Figure 5.**
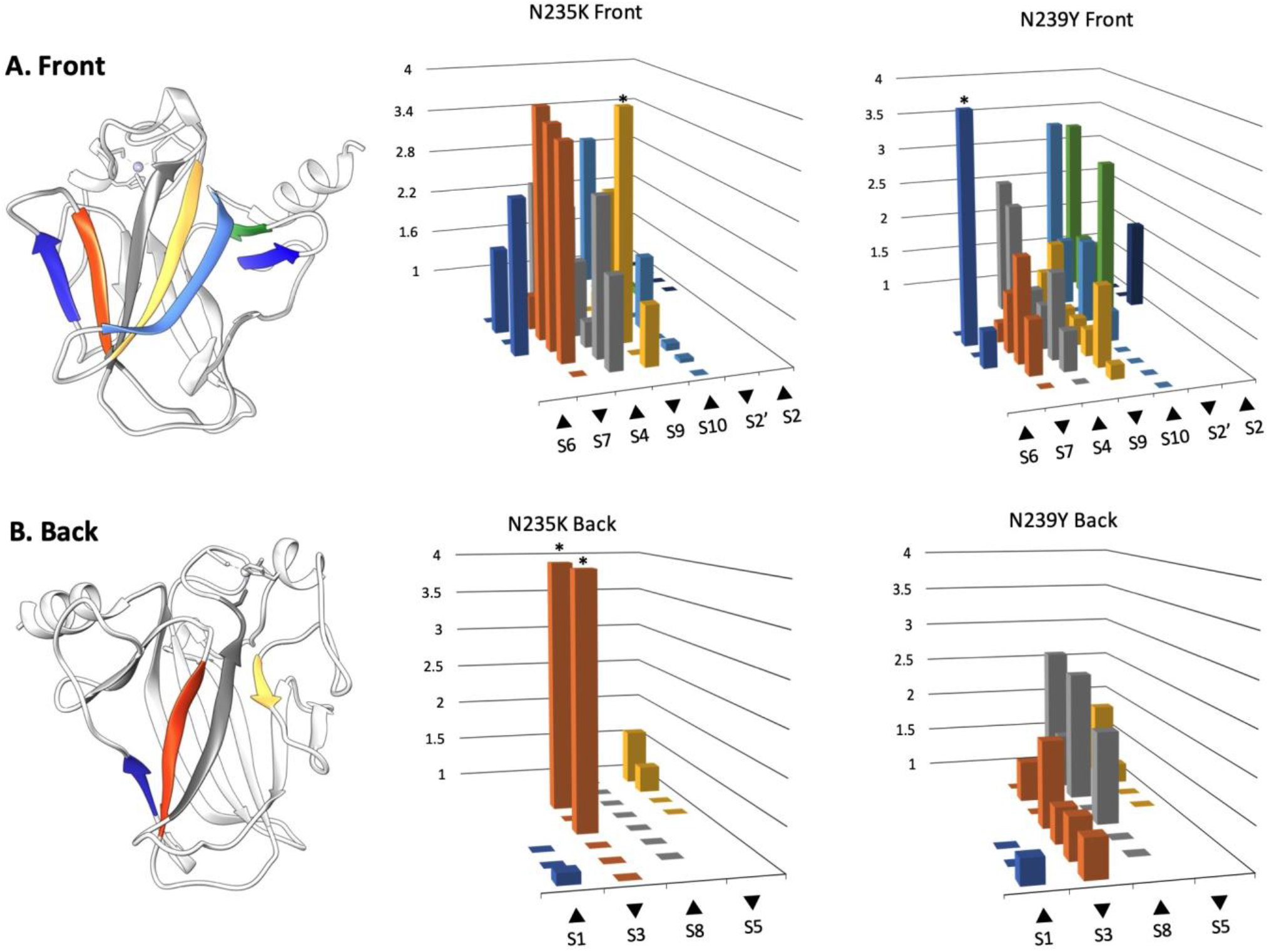
Protection factor changes: mutant/WT ratio. Left panels show the structure of the p53 DBD with β-strands colored to correspond with PF ratios shown in right two panels. Middle and right panels show relative changes in PF as a 3D bar graph representing β-stands organized by their respective position within the p53 protein. Graphs include the β-sandwich ‘front’ side on top **A)** (S6, S7, S4, S9, S10, S2’, S2), and ‘back’ side on bottom **B)** (S1, S3, S8, S5). Bars with asterisks (*) have a mutant/WT ratio above 4. For N235K this includes residue 141(S3), 143(S3), 256 (S9) with ratios of 6.5, 5.4, and 4.6, respectively. For N239Y this includes residue 206 (S6) with a ratio of 30. Larger ratios correlate with a larger increase of PF (slower rate of exchange) in rescue mutants – these positions are stabilized in the mutant compared to WT.

### Relaxation analysis

NMR relaxation studies were performed to determine if these rescue mutants alter backbone dynamics in a fast timescale. Measurement of ^15^N NOE, longitudinal (R1), and transverse (R2) relaxation rates were used to obtain generalized order parameters (S^2^). Relaxation rates for some residues could not be obtained due to rapid signal decay (not enough points to fit) or significant signal overlap. The list of relaxation rates for mutants is calculated are found in **Tables S5 and S6**. We compare these to WT relaxation rates we previously measured (32).

Relaxation rate ratios (R2/R1) are useful for qualitative analysis of dynamics. The usefulness of R2/R1 is the ability to distinguish between fast (ps-ns) and slow (μs-ms) dynamics. Residues with R2/R1 greater than the limit of one standard deviation indicate a region of slow (μs-ms) motions and those less than the limit of one standard deviation are fast (ps-ns) dynamics **(Figure S8)**. We see that both p53 variants have some degree of slow dynamics found in loop region between S3-S4 (residues 147-155), L6, and on S6 **(Figure S8)**. In addition, fast dynamics are observed in loop 1 (L1), loop region immediately presiding S8, and, as expected, in the C-terminus. WT and rescue mutants share fast dynamic timescales in similar locations (32).

Order parameters allow for a more quantitative comparison of flexibility. *Modelfree* uses heteronuclear NOE, R1, R2 measurements and the protein structure to determine backbone flexibility (21, 22). Using the *quadric_diffusion* program (33)(24) we found that the best fitted diffusion tensor model for all variants was an axial symmetry model. N235K and N239Y have an axially symmetric tensor (D _║/⊥_) of, 0.52 and, 0.40 respectively. The average S2 values were 0.85 for N235K and 0.87 for N239Y with correlation times (τ_m_) of 14.1 ns for N235K, and 13.3 ns for N239Y. A plot of calculated S^2^ shows that the most flexible parts are in loop regions **(Figure 6)**. Some flexibility is conserved between rescue mutants; however, other regions show distinct flexibility. For instance, we see similar flexibility in S2’-S3, L6, and S9-S10 loop yet we observe a difference in the later part of L2. To highlight the similarities and differences between WT and rescue mutants we plotted the S^2^ differences between rescue mutants and WT (**Figure 6**). We observe that rescue mutant N235K has regions of increase and decrease flexibility when compared to WT. In contrast, N239Y is predominantly more rigid than WT. For example, we see that the second half of L2 is more rigid in N239Y but more flexible in N235K (**Figure 6**). Both rescue mutants impose distinct changes to backbone flexibility at this time scale.

**Figure 6.**
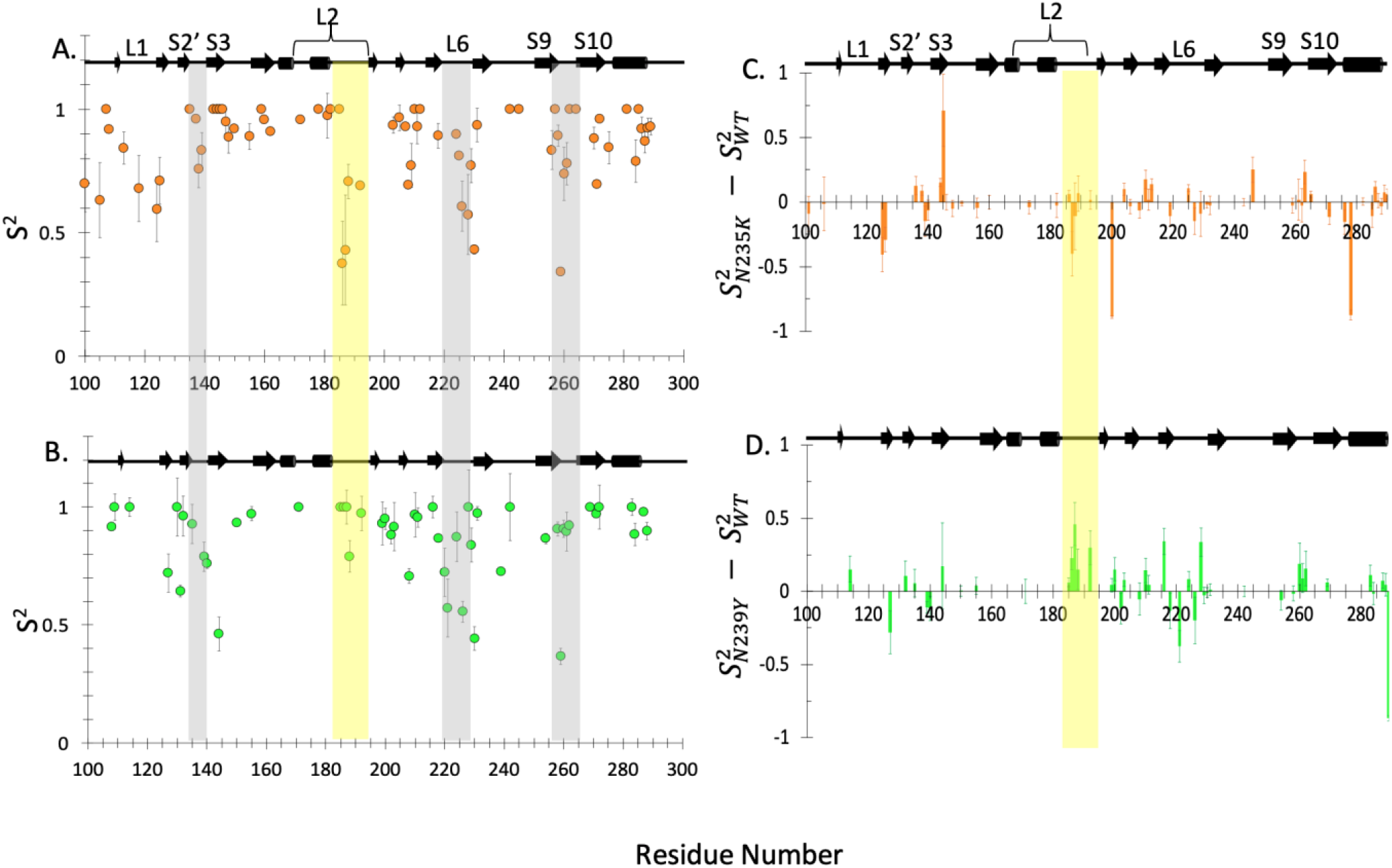
Quantitative characterization of fast dynamics of rescue mutants. Generalized order parameter (S^2^) **A)** N235K, and **B)** N239Y obtained via *Modelfree* analysis (21-23) of relaxation rates R1, R2, and ^1^H-^15^N NOEs at 800 MHz. Similarities between rescue mutant loop dynamics are highlighted with grey vertical boxes. Differences between rescue mutant loop dynamics are highlighted with a yellow vertical box. Differences in order parameters between rescue mutants and WT: **C)** Difference between N235K and WT. **D)** Difference between N239Y and WT.

### Natural occurrence of 235K and 239Y

In order to understand the prevalence of these mutations in nature we performed comparative sequence analysis of p53 sequences in the UniprotKB/Swiss-Prot database. Using the human p53 DBD sequence to search against, we found 3117 sequences in various species. By selecting down to 50% identity, 2670 sequences remained. Asparagines present in the human sequence is also the predominant amino acid in positions 235 (95.92%) and 239 (99.96%). Only one variant at position 239 was found: 239H substitution occurs in *Chanos chanos* (milkfish). In contrast, position 235 can tolerate a range of substitutions including: A, K, R, S, and T. Several species of fish have a conserved a serine (S) at 235 **(Figure 7)**. Eleven species contained a K or R in their respective 235 position; most of these were mouse or rat species.

**Figure 7.**
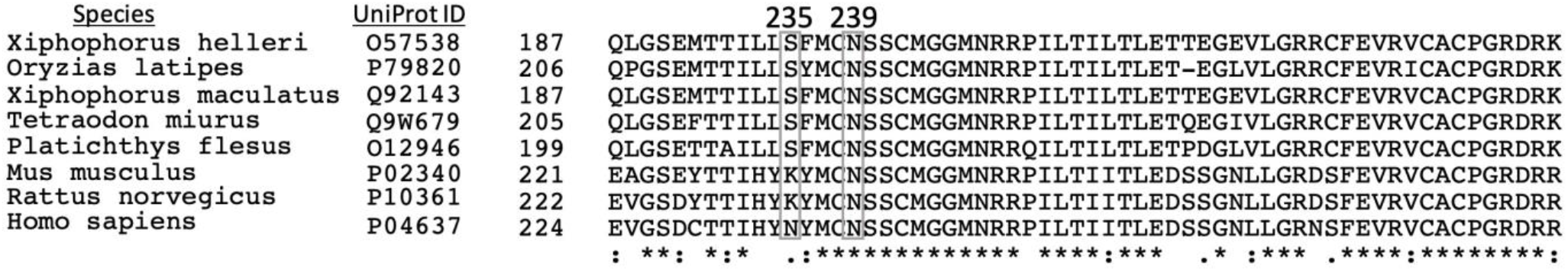
Comparative sequence analysis of p53 across selected species. Protein sequence alignment in the region around 235/239 of the human p53 DBD is shown. Both rat and mouse contain a lysine (K) in their respective position 235. Other species also show variations at position 235; Lys, Arg, Ser, Thr, and Ala are found at 235 in various species. All species have a conserved asparagine (N) in position 239.

## Discussion

Mutations in the DBD can disrupt protein function by either disrupting structure, thermal stability, or DNA binding. N235K and N239Y can rescue the function of multiple cancer mutants (some shown in **Figure 4)**. As previously discovered, rescue of p53 can be accomplished by introducing N239Y and N235K alone (4, 34-36) or in combination with other mutations (4, 34, 35). Both of these rescue mutations maintain WT structure and enhanced DNA binding (5, 9) but prevent the unfolding of some cancer mutants compared to WT. Interestingly, the cancer mutations rescued are spread throughout the DBD, in some cases far from the site of the rescue mutation (**Figure 4**). Here we show that rescue mutations N235K and N239Y dampen motions that lead to hydrogen-deuterium exchange in specific regions throughout the DBD, including the β-sandwich. We also show that these rescue mutants change the dynamics of several loops. The change in dynamics of critical regions of the DBD helps us understand the rescue mechanisms of N235K and N239Y.

In their paper describing the NMR structure of the WT p53 DBD, Canadillas, et al also assessed the backbone hydrogen-deuterium exchange of a few residues. They reported exchange curves for five residues: Cys-238, Thr-253, Thr-256, Tyr-234, and Tyr-236 [supplemental figure 9 in (37)] and calculated the PF for a single residue: Cys238 PF = 7.5 × 10^5^. Our PF for Cys-238 is 5.8 × 10^5^, very close to the published value. Although Canadillas, et al did not calculate PFs for any other positions, we find that our exchange curves are within error for those shown in supplemental figure 9 of (37).

### Protection in the β-sandwich scaffold

Past research on N235K and N239Y indicate that both rescue mutations have far-reaching effects within the DBD. The mechanism by which N235K restores stability and function has been explored through molecular simulations (8) and experimental methods (9). Both studies attribute the increase in stability to newly formed salt bridges created by the lysine substitution. Wallentine and coworkers speculate that this newly formed salt bridge is sufficient to stabilize the entire β-sandwich (9). Our data corroborates this speculation of far-reaching stability. We found that 35 amides were more protected in N235K than in the WT; most of which are spread through the β-sandwich including residues 156, 206, 258, 259, 268, 270, which are located 15-23 Å from the site of rescue mutation. Eighteen of these amides have a PF increase greater than two-fold **(Figure 4)** compared to WT. Interestingly, the highly protected amides in N235K are located on one side of the β-sandwich scaffold in strands 4, 6, 7, 9, and 10 **(Figure 5)**, opposite to the site of mutation. In contrast, the regions that are destabilized by the rescue mutation are found in proximity to strand 3 and 8, on the same side of the β-sandwich as the mutation. Thus, N235K acts as a fulcrum in increasing rigidity in one sheet, but slightly destabilizing the other. The balance of increased protection in one region and decreased in another is consistent with only a very minor increase in thermal stability observed in N235K compared to WT (8, 9). In contrast, the increase in stability in N239Y was uniform on both sides of the β-sandwich. The effects of N239Y are widespread; 45 amides had larger protection in N239Y compare to WT; 18 had a two-fold increase in protection. These 18 highly protected amides are widely spread throughout both sides of the β-sandwich scaffold **(Figure 5)**. This extensive protection indicates that the hydrophobic core is well stabilized and provides a rationale for the increase in thermal stability of ∼ 1 kcal mol^-1^ observed in N239Y (5, 7, 9).

N235K and N239Y are both able to rescue cancer mutant V157 even though position 157 is 15 Å away from position 235, and 24 Å away from position 239 (157 is shown in **Figure 4**). V157F is characterized as one of the strongest β-sandwich missense mutations since it destabilizes the DBD by 3.6 kcal mol^-1^ (6, 8, 38). It has been proposed that the substitutions of a small (valine) to the large (phenylalanine) hydrophobic residue disrupts hydrophobic packing of the DBD (38). Our data shows that both N235K and N239Y stabilize the distant region where 157 is localized. In N235K amide 157 has a three-fold increase in PF compared to WT. In addition, 157 is found in the region of the protein that shows a clear trend of increased stability, including the surrounding strands 4, 7, and 9 **(Figure 5)**. A similar trend is observed in rescue mutant N239Y. The increased rigidity in this region of the DBD counteracts the destabilization caused by V157F, allowing for the restoration of function and stability. This stabilizing effect also likely to be true for other oncogenic mutations found in the same region. Danziger and coworker found seven cancer mutants that can be rescued by N235K; these include: C141Y, V157F, R158L, Y205C, Y220C, V272M, and E286K (35). The effect of rescue mutations on cancer mutants at positions 157, 158 were also confirmed in other studies (9, 36). Five of the seven rescuable cancer mutants are found on the β-sheet of N235K that we find to be highly protected. Position 141 is also one of the cancer mutants rescued by N235K and we find it to be one of the most protected residues in N235K and with significantly increased protection compared to WT. Our hydrogen exchange results show a strong correlation between positions with increased protection from hydrogen exchange that are predictive of distant cancer mutants rescued.

### Stabilization of the aggregation region

WT p53 DBD is inherently prone to aggregation and amyloid formation compared to other proteins of the same family (39). These aggregation tendencies have also been shown to be increased in cancer mutations of p53 (40, 41). Several investigators have attributed this aggregation to transient exposure of sidechains and backbone hydrogen bonds of residues 251-257 in strand 9 (S9) (39, 41, 42). Our results show that both rescue mutations significantly increase the stability of backbone amide hydrogen bonding in the 251-257 region of S9. N235K had an increase in protection for all residues in S9 compared to WT, except for residues 253 and 257 that were slightly less than that of WT. In N239Y all residues in this region had an increase in protection, with the largest increase in protection observed for residues 253 and 257. Thus, the pattern of protection in S9 is unique to each rescue mutation. In combination, these rescue mutations could increase the protection of the entire S9 strand. N235K and N239Y together would impose an additive protective effect that could explain the synergistic stabilization previously reported in the global rescue motifs involving both of these rescue mutations (4, 7, 9).

### Protection in the DNA-binding interface: S2/S2’/S3/S10 strands, L2/L3, Zn binding)

The interactions caused by N239Y induce protection in the structures that support the DNA binding interface. Wallentine and coworkers identified novel hydrophobic interactions created by the newly introduced 239 tyrosine and the native 137 leucine (9). Our analysis of N239Y shows that both of the strands S2’ (residues 132-135), and S3 (residues 141-146) that surround 137 have an increase in protection factor for all positions measured. In addition, residues 127 (S2), 135 (S2’), 236 (S8), 238 (L3), 273 (S10), and 275 (S10) have PFs that are two-fold or greater compare to WT, which indicate high stability for these positions near the DNA binding interface.

The stability caused by the novel hydrophobic interaction in N239Y may also play a role in stabilizing the Zn binding region, which includes loop 2 (L2) and loop 3 (L3). Zinc binding is important in maintaining structure stability in the p53 DBD (43, 44). The binding of Zn is coordinated by residues 176 and 179 in Loop 2, and residues 238 and 242 in Loop 3. When zinc is absent loops 2 and 3 dissociate, which gives rise to an inactive p53 structure that is prone to aggregation (43). N235K and N239Y have been shown to restore function to several cancer mutations localized in the Zn binding region of L2 and L3, including G245C, G245S, V173L, and V173M for N239Y, and G245S, V173L for N235K (4, 7, 15, 34). Our data supports the hypothesis that the L2/L3 interface is more stable in both rescue mutants. Both mutants contain L2 and L3 amides that experience an increase in protection. N235K has an increase in protection for 163, 237, 238. Position 237 in N235K had the largest PF observed in all residues analyzed and the sidechain of 238 is directly involved in Zn binding. N239Y has a significant increase in protection for residues 163, 193 and 238. Notably, amide 193 is only protected in the rescue mutants but not in the WT. Furthermore, backbone amides at positions 163 and 238 are buried in the L2/L3 loop region with a solvent accessible surface area of 0 Å^2^ for 238 and 4 Å^2^ for 163 and these are not involved in backbone hydrogen bonding. The increase in protection can only be attributed to the structure protection that arise from a well-stabilized L2/L3 loop structure.

Further evidence of increased stability in the DNA binding interface comes from the increase rigidity observed in S^2^ values of N239Y in the latter half of L2, which spans residue 183-194 (**Figure 6**). Principle component analyses of residue positions in multiple p53 DBD crystal structures have shown this part of L2 to be highly flexible (45). S^2^ calculations of WT show that this flexibility is confirmed to be present in WT (32) and N235K but not N239Y (**Figure 6**). Previous studies have shown that the hot-spot cancer mutant G245S increase flexibility in this region, specifically between residues 191-193 (13). N239Y rescues function for G245S; the increase in rigidity in L2 may be the main mechanism of rescue for this particular cancer mutant. Together, the increased protection seen in the DNA binding interface and specific rigidity in residue 183-194 of N239Y may be critical to the mechanism of rescue for N239Y.

### Loop dynamics: Loop 1 (113-123), Loop 6 (221-230)

Protein flexibility is intimately related to protein function. L1 has been shown to be involved in dynamic DNA binding (37). Principle component analysis (45), molecular dynamics (46, 47), and NMR studies (37) of p53 DBD have shown L1(113-123) to be highly flexible. This flexibility is observed in WT (32) and N235K (**Figure 6**). NMR studies of WT p53 DBD suggested L1 has a slow motion (μs-ms) (37). We find that the flexibility in L1 is more predominant in N235K compared to WT S^2^ values (32) (**Figure 6**). The increase in flexibility is consistent with local changes in chemical shifts for residue 117 in N235K compared to WT (**Figure 2**). As suggested previously, the dynamics of L1 have been proposed to play a role in the transcription factor mechanism of tread-milling to find DNA targets (46). Although N235K reverses the destabilizing effects of some cancer mutants, it shows an increase in flexibility in L1 that could affect DNA binding.

Residues 221-230 (referred to as L6 in (47)) has been shown to be flexible in the DBD (26, 47). This flexibility is apparent in crystal structure B-factors for WT, N235K, and N239Y (**Figure S9**). All three variants have an increase in B-factors between residues 210-230, with WT having the largest B-factors between residues 224-228. WT (32) and rescue mutant S^2^ values also show this loop to be the most flexible region in all p53 variants (**Figure 6**). Of all the loop regions, L6 seem to have the most common dynamics across all three p53 variants and is not affected by the rescue mutations studied here.

### Enhanced flexibility in N235K and N239Y: S2’-S3 loop (136-140), S9-S10 loop (259-263)

Both rescue mutants show increased flexibility in regions not previously observed in WT (32). S^2^ values indicate that both rescue mutants have increased flexibility in S2’-S3 loop (residue 136-140) and in the S9-S10 loop (259-263) (**Figure 6**). These rescue mutants are in close proximity to the S2’-S3 loop in the tertiary structure of the DBD. This is highlighted by the increase in chemical shift changes within the S2’-S3 loop induced by both rescue mutants (**Figure 2**). The proximity of this loop to the rescue mutants provides a likely explanation for the change in flexibility. Interestingly, the S^2^ plots show that S2’-S3 loop of N239Y is more flexible than in the WT (32); however as discussed above, we found slower rates of hydrogen exchange for the same region. From both of these results we can conclude that this part of the DNA binding interface is resistant to unfolding but the loop has increased motions in the fast timescale in N239Y.

In contrast, the S9-S10 loop is far from the sites of mutation (> 24.6 Å). Chemical shift analysis does not indicate any changes to local structure within the S9-S10 loop (**Figure 2**). The increased flexibility seen in S^2^ values in this distant location serves as evidence of the far-reaching effects that both N235K and N239Y impose on the DBD.

### Sequence comparisons

The p53 sequences of both mouse and humans have been studied and compared. They are very similar in sequence (∼ 96.9% similarity) and structures (RSMD < 0.6 Å^2^). Khoo and coworkers measured and compared the stability of p53 DBD across a wide variety of species and concluded that p53 evolved to be less stable in humans (48, 49). The fact that tyrosine is never found as a substitute at position 239 in any species may be related to the requirement that p53 be degraded easily under certain cellular conditions. N239Y is strongly stabilizing and would limit the ability for cells to regulate p53 function.

The substitution of arginine to lysine at position 235 may have played a critical role in the evolution of p53 from a more stable form in some species to less stable in humans. Our sequence analysis revealed that both mice and rats contain a lysine in the 235 position. Consequently, the stabilizing groups within the p53 fold are distributed differently in these animals compared to human p53. Since 235K is known to be more resistant to cancer mutations, animal studies of cancer using mice or rats should include the humanized p53 sequence.

Ten other positions are found to be conserved between rat and mice but different in humans. These includes positions 110, 123, 133, 165, 185, 202, 210, 229 268, 289. Out of these ten positions, three have been discovered to have stabilizing effects. These include L133, K235, D268. (4, 5, 34, 36). Interestingly, the N268D rescue mutation was discovered through sequence comparisons across species (5, 50). Given that p53s in other species are more stable, sequence comparisons could guide the discovery of future rescue mutations.

## Conclusions

In summary we found that both N235K and N239Y have far reaching effects on the dynamics of the DBD. These rescue mutants increase overall folding stability of the DBD as measured by hydrogen exchange, while at the same time modify dynamics at specific loop regions. N235K has an asymmetric stabilizing effect on the β-sandwich with one sheet stabilized, while imposing enhanced flexibility in the S2’-S3 and S9-10 loops. N239Y has a stabilizing effect on both sheets of the β-sandwich, reduces flexibility on L2, while imposing the same flexibility in S2’-S3 and S9-10 loops. The change in dynamics imposed by these rescue mutants should play a critical role in their ability to rescue cancer mutants. Therapeutic interventions can be guided by pursuing drugs that can recapitulate the dynamics of these rescue mutants. Stabilization of residues by slowing the rate of hydrogen exchange may serve as a metric in the development of anticancer drugs to block p53 unfolding and aggregation.

## Author Contributions

**JS:** Data analysis, Preparation of figures, Writing - Preparation of Original Draft; **CM:** Performed experiments, Data analysis, Writing - Review & Editing; **AA:** Data Analysis;**MS-B:** Protein purification, Writing - Review & Editing; **DM:** Protein purification; **MJC:** Conceptualization, Data Curation, Supervision, Writing - Review & Editing

## Acknowledgements

Funding was provided by the California Cancer Research Coordinating Committee (CRCC9-550862-36240) and NIH-IMSD training grant GM055246.

## Supplemental Information

**Figure S1.**
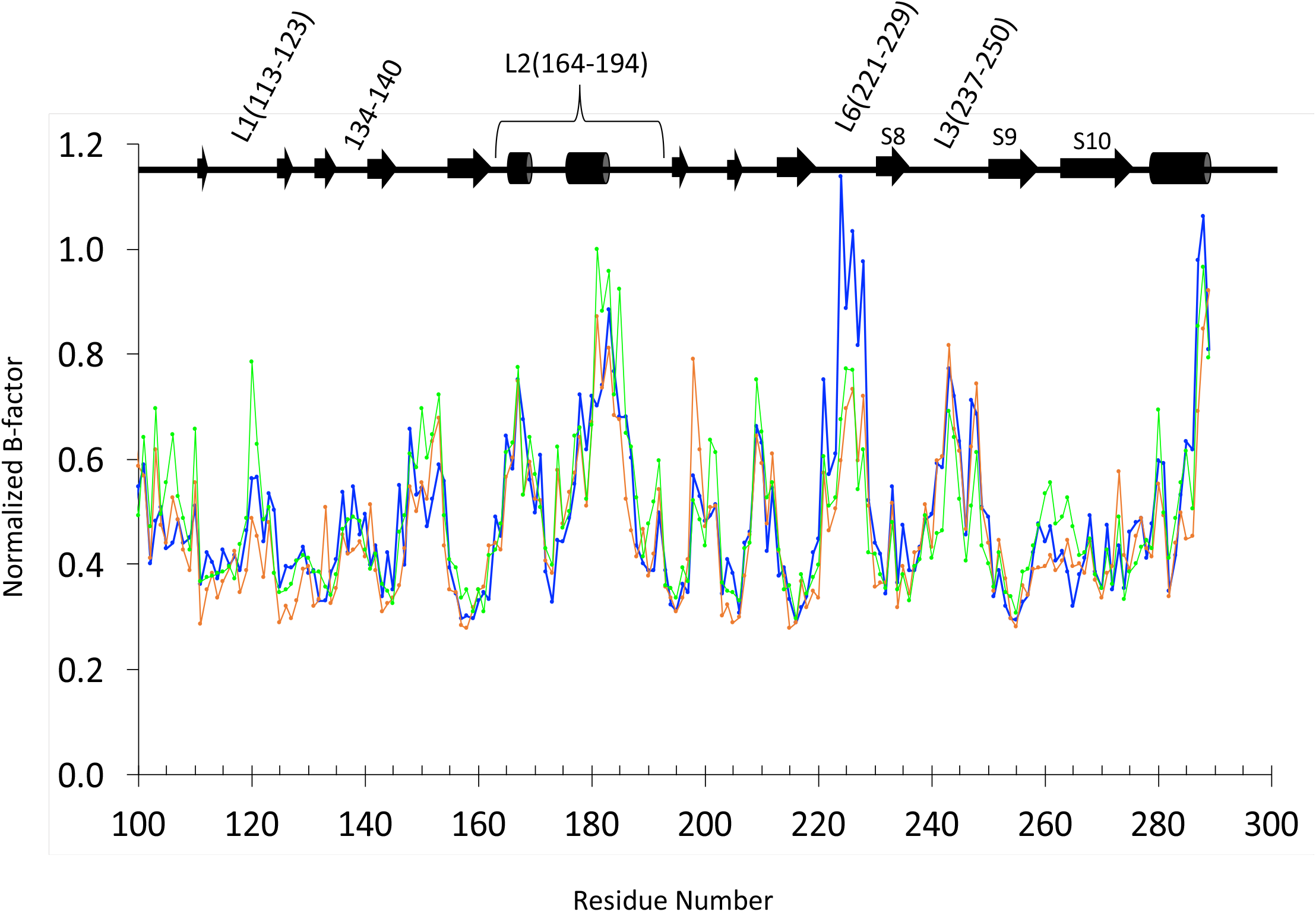
Normalized B-factors from crystal structures of WT (PDB: **<u>2XWR</u>**, blue), N235K (PDB: **<u>4LO9</u>**, orange), N239Y (PDB: **<u>4LOE</u>**, green). B-factor trends are in agreement between all p53 variants with increases in B-factors localized to loop regions.

**Figure S2.**
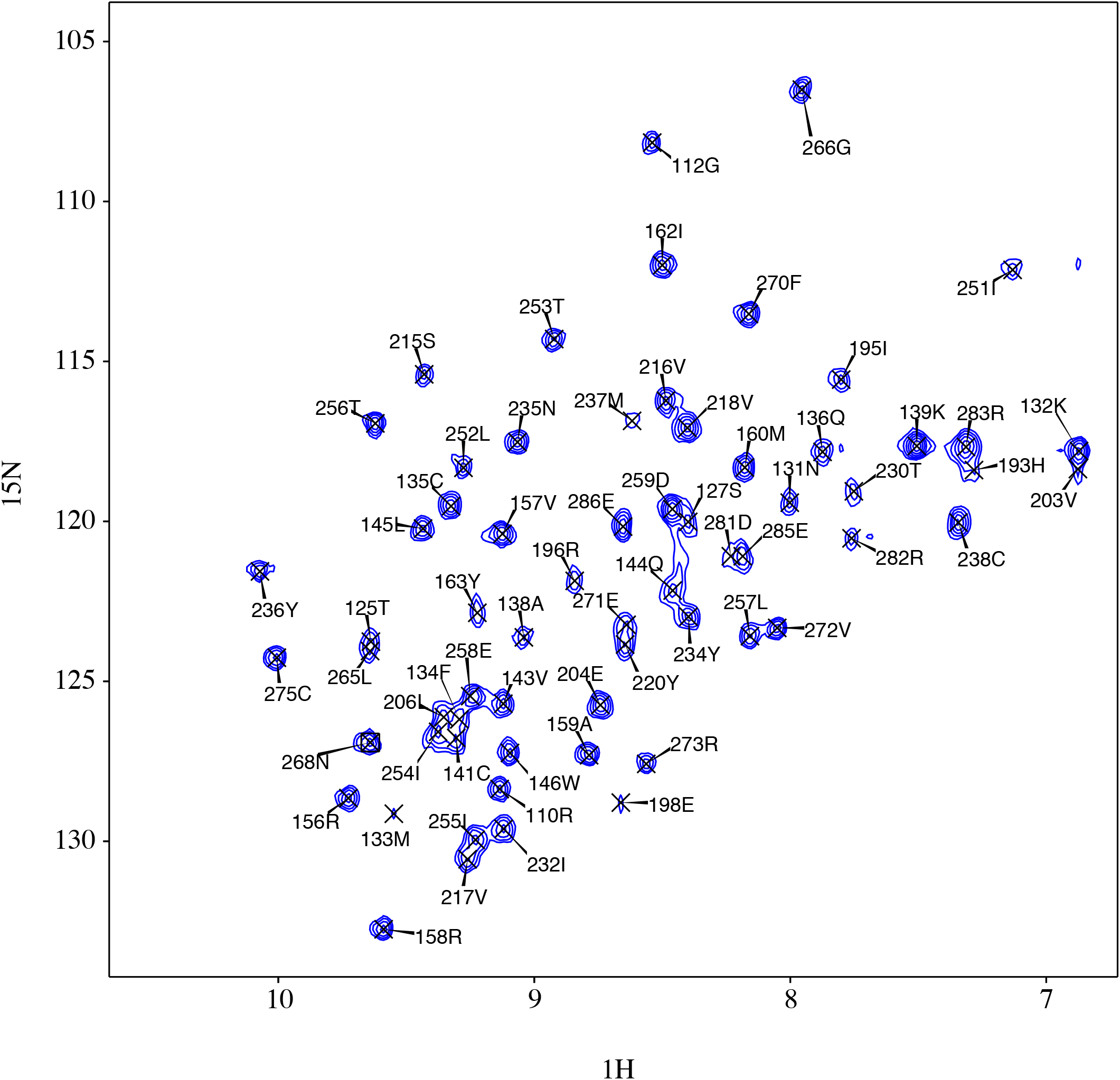
Peak assignments of the first time point after initiation of exchange with D_2_O for WT p53 (16 minutes).

**Figure S3.**
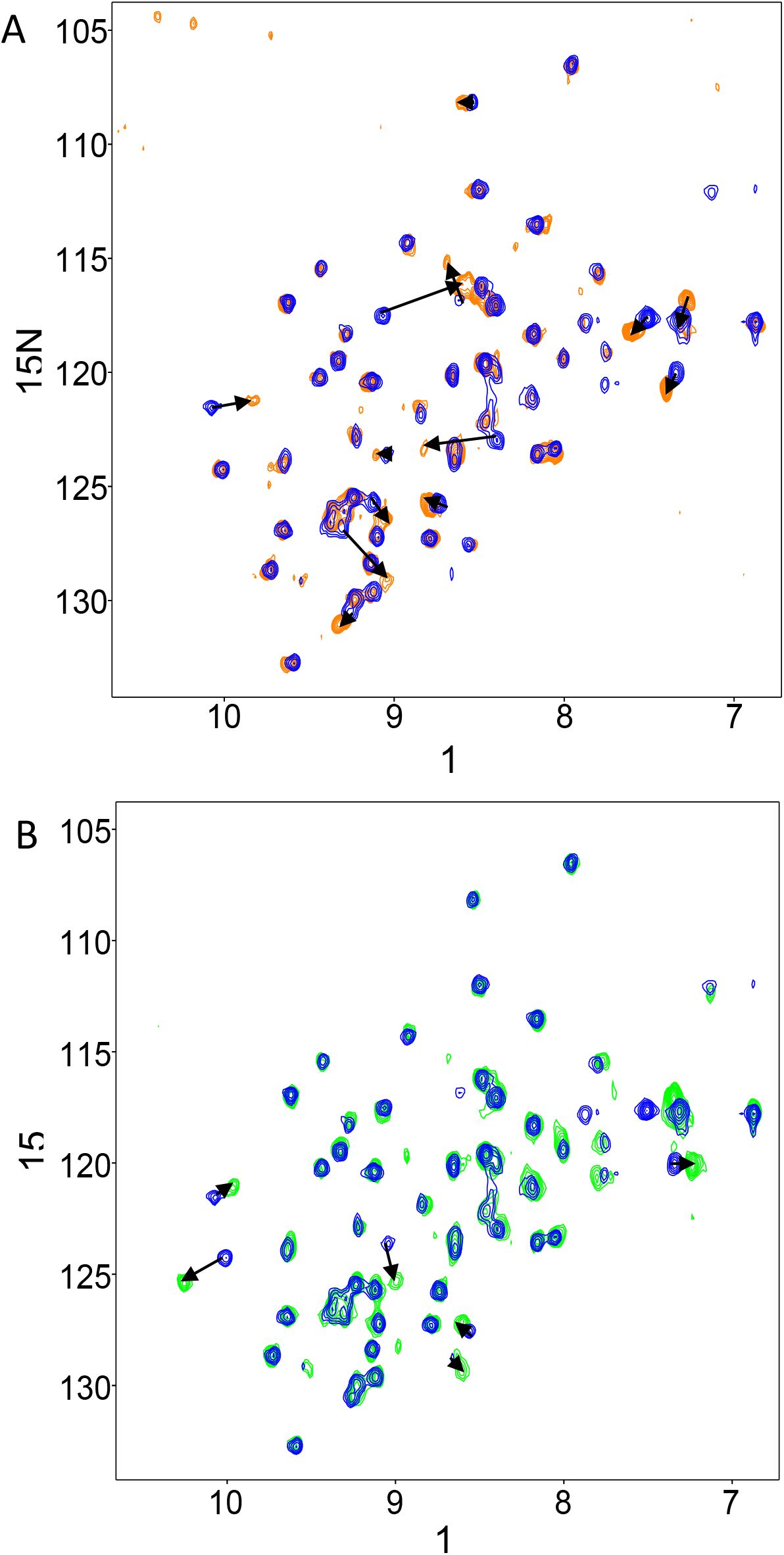
Overlay of WT spectra with **(A)** N235K (orange) and **(B)** N239Y (green). Spectra of the first time point after initiation of exchange. Shifted peaks were assigned according to backbone NOEs.

**Figure S4.**
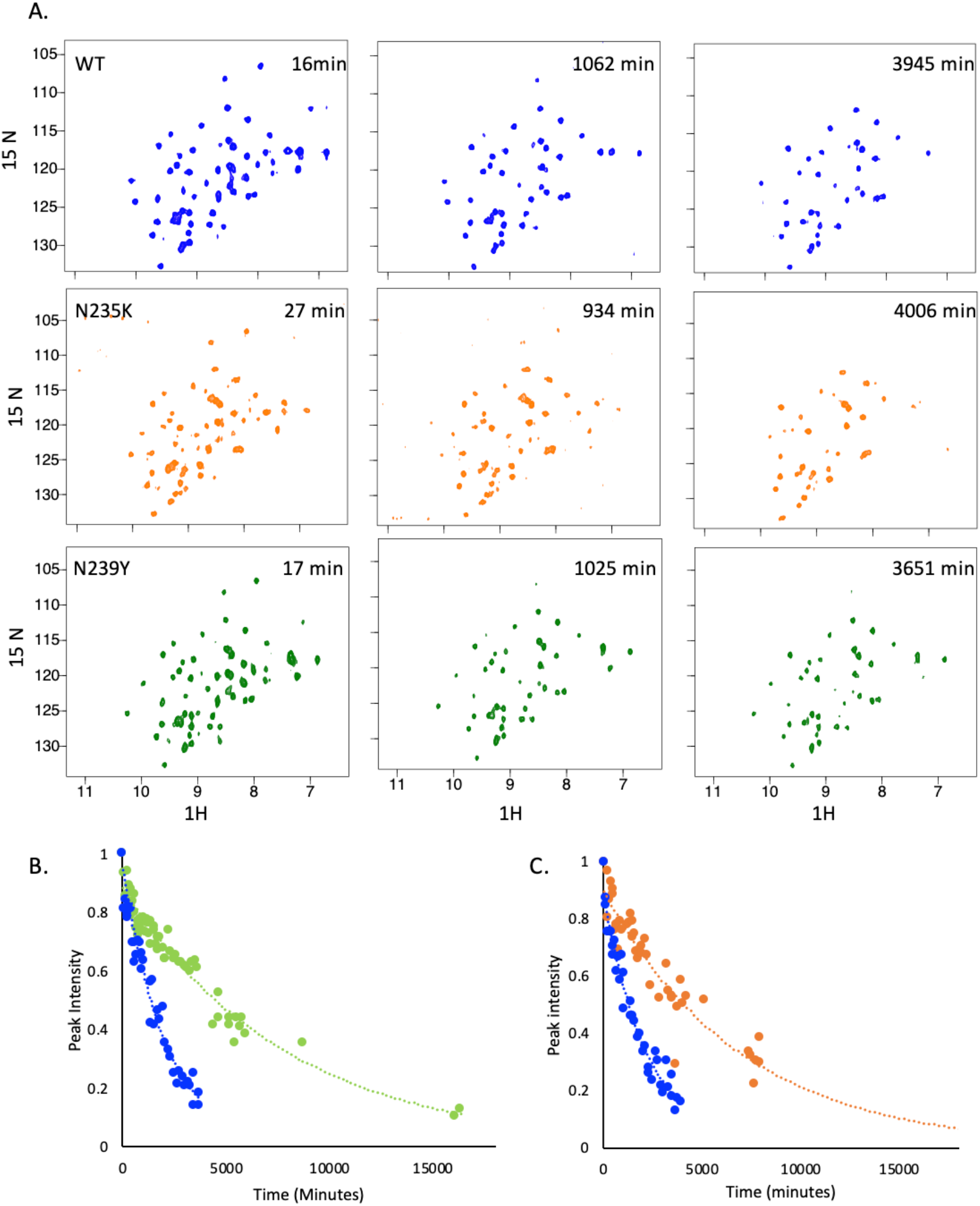
**(A)** ^15^ N-HSQC spectra of p53 wild-type (blue), N235K (orange), and N239Y (green) at different time points after transferred into D_2_O. Peaks were integrated at each time point over a period of two weeks and plotted as a function of time. Hydrogen exchange rates (k_ex_) were calculated from a fit to a single exponential. Representative data: (**B)** 135Cys hydrogen signal intensities in N239Y (green) and WT (blue) p53 DBD. **(C)** 204Glu hydrogen signal intensities in N235K (orange) and WT (blue) p53 DBD.

**Figure S5.**
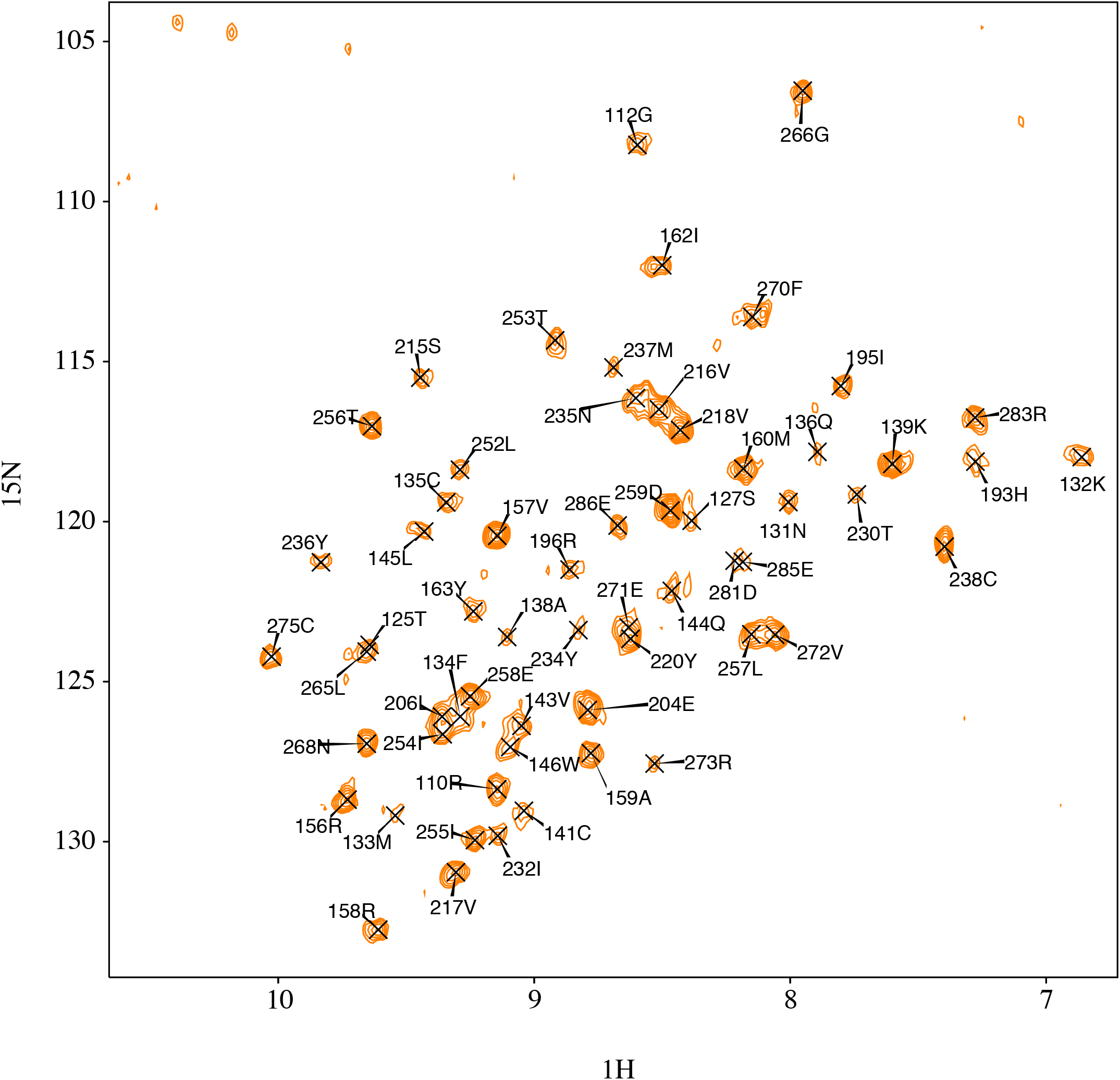
Peak assignment for rescue mutants N235K. Spectra at 27 minutes after initiation of exchange with D_2_O.

**Figure S6.**
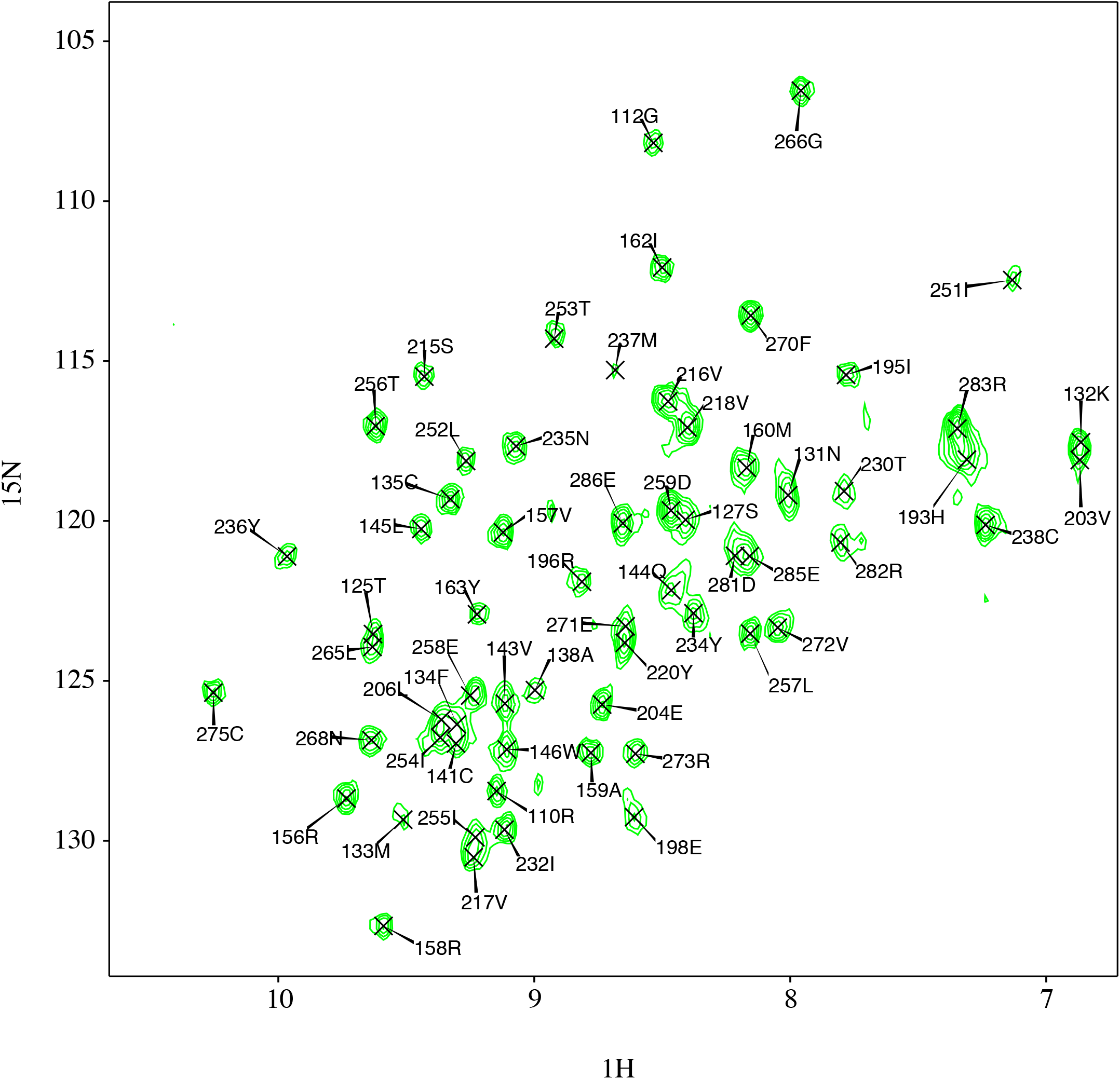
Peak assignment for rescue mutants N239Y. Spectra at 17 minutes after initiation of exchange with D_2_O.

**Figure S7.**
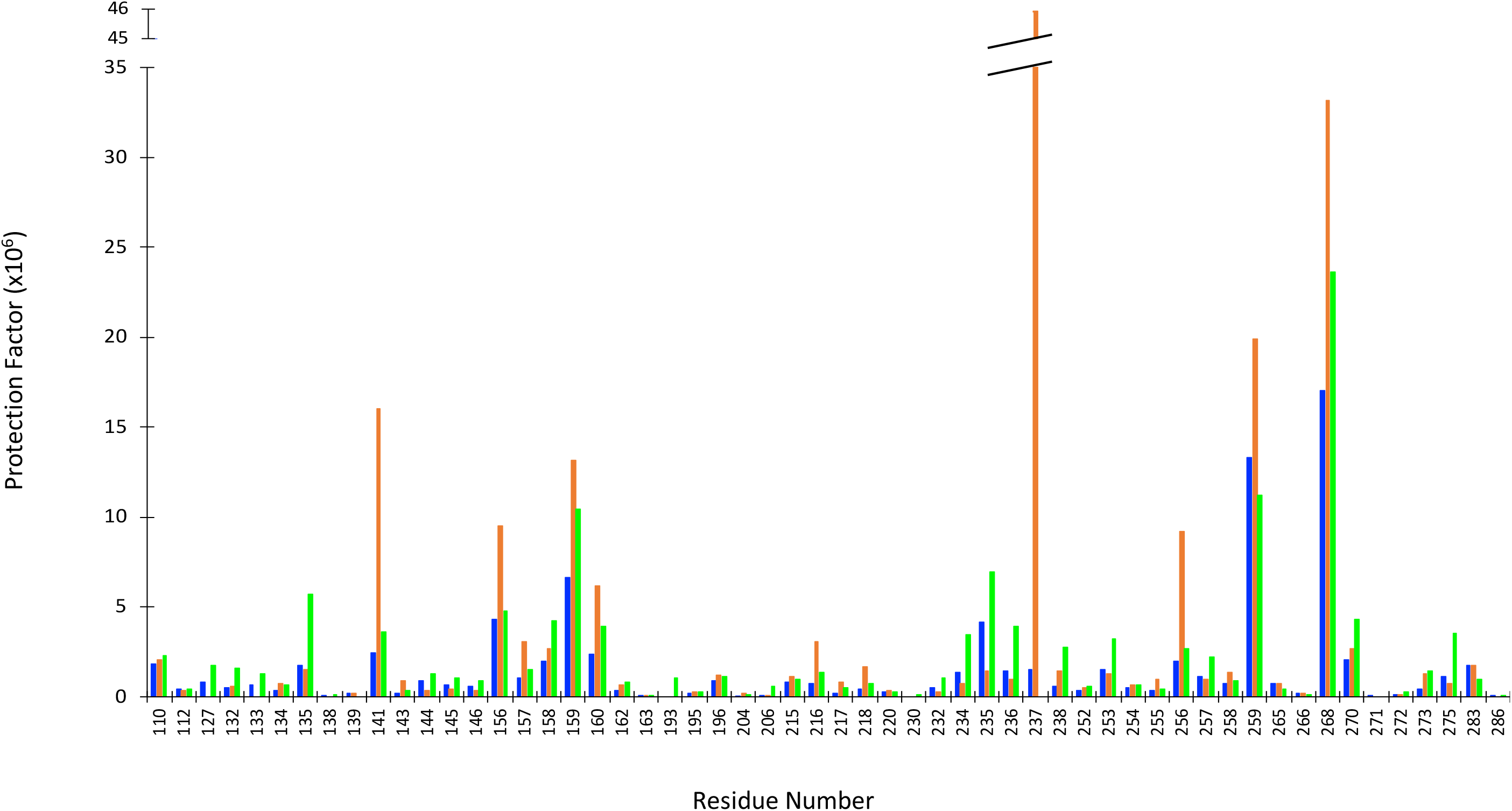
Protection factors of WT and rescue mutants. 35 amide hydrogens had an increase in protection factors in the rescue mutant N235K (orange) compared to WT p53 (blue). A total of 45 amide hydrogens have an increase in protection factors in the rescued mutant N239Y (green) compared to WT. Residues 193 and 230 are protected in N239Y but exchange too fast to be detected in WT or N235K.

**Figure S8.**
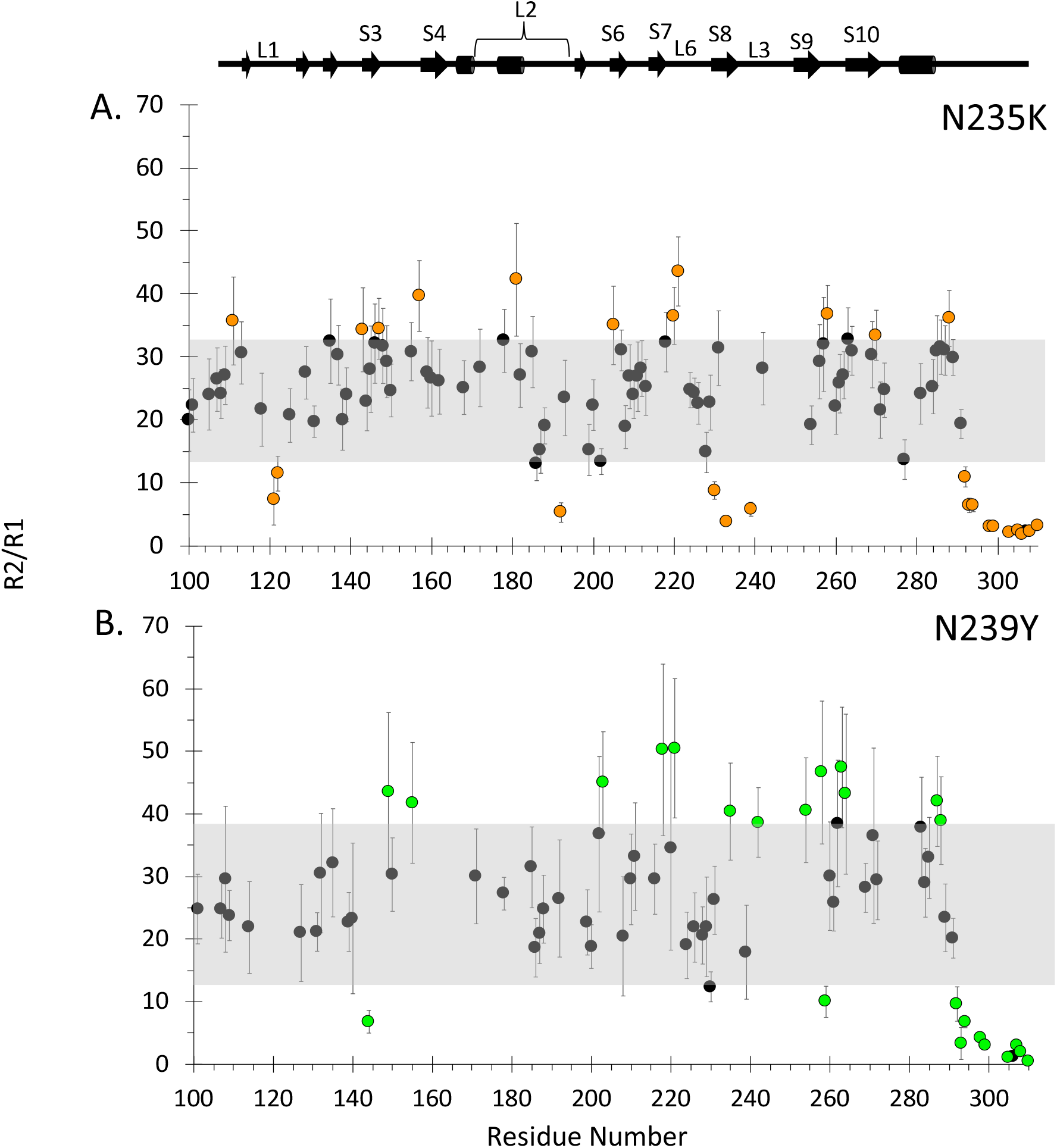
R2/R1 plot for qualitative analysis of back bone dynamics in **(A)** N235K, and **(B)** N239Y. Grey rectangles highlight mean values ± 1 SD of all R2/R1 for each p53 variant. Colored data points highlight the residues that are outside the mean ± 1 SD. Missing points in N239Y reflect signals that decayed too fast to reliably calculate a rate.

**Figure S9.**
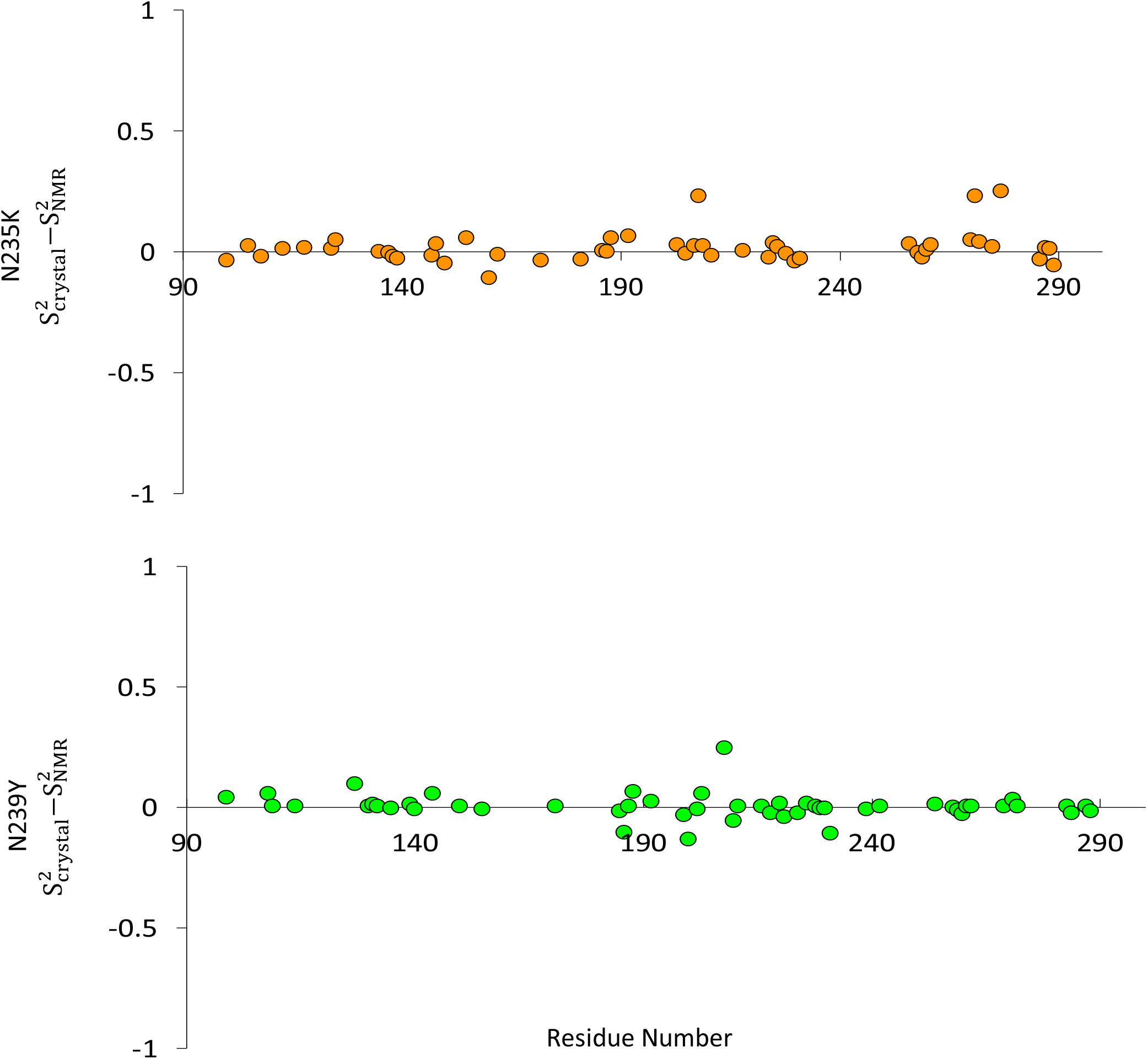
Comparison of NMR S2 Calculated from Mutant Crystal or WT NMR Structures. Use of mutant p53 crystal or WT NMR coordinate files do not change *Modelfree* S^2^ calculation results. **(A)** Difference between S^2^ values calculated from N235K relaxation data when using N235K crystal (PDB: **<u>4LO9</u>**) or WT NMR as the structure file (PDB: **<u>2FEJ</u>**) **(B)** Difference in S^2^ values calculated from N239Y data when using N239Y crystal (PDB: **<u>4LOE</u>**) or WT NMR as the structure file (PDB: **<u>2FEJ</u>**).

**Table S1.** Chemical Shifts of residues that showed significant change in peak positions compared to WT. Indicates residue that did not show significant chemical shift in that respective rescue mutants.

| Residue # | WT |  | N235K |  | N239Y |  |
| --- | --- | --- | --- | --- | --- | --- |
|  | H position (ppm) | <sup>15</sup>N position (ppm) | H position (ppm) | <sup>15</sup>N position (ppm) | H position (ppm) | <sup>15</sup>N position (ppm) |
| 117 | 8.67 | 111.24 | 8.65 | 111.76 | - | - |
| 124 | 7.48 | 120.32 | 7.51 | 119.99 | - | - |
| 125 | 9.63 | 123.66 | 9.59 | 123.05 | - | - |
| 126 | 8.72 | 128.94 | 8.77 | 129.24 | - | - |
| 137 | 8.61 | 125.53 | 8.50 | 125.60 | - | - |
| 138 | 9.03 | 123.57 | - | - | 9.05 | 123.03 |
| 139 | 7.50 | 117.77 | 7.59 | 118.21 | 7.35 | 117.14 |
| 141 | 9.30 | 126.79 | 9.09 | 129.57 | - | - |
| 143 | 9.11 | 125.70 | 9.02 | 126.38 | - | - |
| 175 | 8.00 | 116.30 | 8.00 | 116.63 | - | - |
| 180 | 8.83 | 120.35 | 8.85 | 120.07 | - | - |
| 186 | 8.18 | 124.78 | 8.23 | 124.48 | - | - |
| 187 | 8.54 | 109.33 | 8.52 | 109.62 | - | - |
| 196 | 8.84 | 121.94 | 8.85 | 121.55 | - | - |
| 199 | 8.78 | 110.83 | 8.77 | 111.56 | - | - |
| 200 | 8.22 | 117.32 | 8.22 | 117.60 | - | - |
| 203 | 6.87 | 118.33 | 6.90 | 118.89 | - | - |
| 217 | 9.24 | 130.54 | 9.28 | 130.86 | - | - |
| 234 | 8.38 | 122.94 | 8.74 | 123.45 | - | - |
| 235 | 9.06 | 117.56 | 8.56 | 115.98 | - | - |
| 236 | 10.07 | 121.65 | 9.85 | 121.37 | 9.97 | 121.13 |
| 237 | 8.60 | 116.83 | 8.58 | 115.16 | 8.60 | 115.72 |
| 238 | 7.33 | 120.27 | 7.37 | 120.77 | - | - |
| 239 | 8.45 | 118.33 | 8.41 | 118.04 | 8.45 | 117.88 |
| 240 | 9.60 | 120.16 | - | - | 9.63 | 119.64 |
| 241 | 8.13 | 109.49 | - | - | 7.96 | 108.95 |
| 242 | 7.80 | 122.18 | - | - | 7.91 | 122.29 |
| 273 | 8.55 | 127.70 | - | - | 8.60 | 127.36 |
| 275 | 9.99 | 124.27 | - | - | 10.25 | 125.45 |
| 277 | 8.74 | 115.75 | - | - | 8.83 | 115.88 |
| 279 | 8.89 | 104.80 | - | - | 8.92 | 105.25 |

**Table S2.** Protection Factors for WT p53.

| Residue | $k_{\text{ex}} (10^{-4}) / \text{min}$ | Error | $k_{\text{rc}} (10^{-4}) / \text{min}$ | PF ($10^6$) |
| --- | --- | --- | --- | --- |
| 110Arg | 1.24 | 0.11 | 2.24E+06 | 1.81 |
| 112Gly | 4.38 | 0.30 | 1.86E+06 | 0.43 |
| 127Ser | 5.16 | 0.49 | 4.27E+06 | 0.83 |
| 132Lys | 5.81 | 0.30 | 3.09E+06 | 0.53 |
| 133Met | 2.98 | 1.43 | 2.09E+06 | 0.70 |
| 134Phe | 3.39 | 0.54 | 1.20E+06 | 0.36 |
| 135Cys | 4.48 | 0.19 | 7.76E+06 | 1.73 |
| 138Ala | 26.20 | 5.27 | 1.00E+06 | 0.04 |
| 139Lys | 6.32 | 0.15 | 1.48E+06 | 0.23 |
| 141Cys | 4.37 | 0.31 | 1.07E+07 | 2.45 |
| 143Val | 1.07 | 0.10 | 1.86E+05 | 0.17 |
| 144Gln | 1.50 | 0.15 | 1.35E+06 | 0.90 |
| 145Leu | 0.99 | 0.09 | 6.76E+05 | 0.68 |
| 146Trp | 0.65 | 0.09 | 3.89E+05 | 0.60 |
| 156Arg | 0.71 | 0.08 | 3.09E+06 | 4.34 |
| 157Val | 0.53 | 0.06 | 5.37E+05 | 1.02 |
| 158Arg | 0.70 | 0.09 | 1.41E+06 | 2.02 |
| 159Ala | 0.41 | 0.06 | 2.69E+06 | 6.64 |
| 160Met | 0.67 | 0.08 | 1.59E+06 | 2.38 |
| 162Ile | 0.90 | 0.09 | 3.02E+05 | 0.34 |
| 163Tyr | 37.70 | 3.64 | 5.13E+05 | 0.01 |
| 195Ile | 1.14 | 0.13 | 1.86E+05 | 0.16 |
| 196Arg | 1.27 | 0.16 | 1.15E+06 | 0.91 |
| 204Glu | 4.58 | 0.18 | 3.63E+05 | 0.08 |
| 206Leu | 30.50 | 4.95 | 4.79E+05 | 0.02 |
| 215Ser | 6.60 | 0.51 | 5.25E+06 | 0.79 |
| 216Val | 0.83 | 0.07 | 6.46E+05 | 0.78 |
| 217Val | 1.12 | 0.09 | 2.34E+05 | 0.21 |
| 218Val | 0.53 | 0.05 | 2.34E+05 | 0.45 |
| 220Tyr | 1.65 | 0.10 | 5.01E+05 | 0.30 |
| 232Ile | 0.93 | 0.09 | 4.79E+05 | 0.51 |
| 234Tyr | 0.89 | 0.07 | 1.20E+06 | 1.35 |
| 235Asn | 1.34 | 0.11 | 5.62E+06 | 4.18 |
| 236Tyr | 1.24 | 0.11 | 1.82E+06 | 1.47 |
| 237Met | 1.17 | 0.20 | 1.78E+06 | 1.52 |
| 238Cys | 15.00 | 0.93 | 8.71E+06 | 0.58 |
| 252Leu | 0.70 | 0.11 | 2.51E+05 | 0.36 |
| 253Thr | 0.56 | 0.08 | 8.51E+05 | 1.53 |
| 254Ile | 0.93 | 0.10 | 4.79E+05 | 0.52 |
| 255Ile | 0.54 | 0.08 | 1.78E+05 | 0.33 |
| 256Thr | 0.41 | 0.06 | 8.13E+05 | 2.00 |
| 257Leu | 0.61 | 0.08 | 6.76E+05 | 1.11 |
| 258Glu | 0.41 | 0.06 | 3.09E+05 | 0.75 |
| 259Asp | 1.47 | 0.08 | 1.95E+07 | 13.30 |
| 265Leu | 0.36 | 0.12 | 2.63E+05 | 0.72 |
| 266Gly | 11.00 | 0.56 | 1.86E+06 | 0.17 |
| 268Asn | 0.49 | 0.06 | 8.32E+06 | 17.03 |
| 270Phe | 0.90 | 0.07 | 1.86E+06 | 2.07 |
| 271Glu | 28.60 | 5.38 | 5.76E+05 | 0.02 |
| 272Val | 1.51 | 0.13 | 2.29E+05 | 0.15 |
| 273Arg | 3.36 | 0.19 | 1.41E+06 | 0.42 |
| 275Cys | 4.26 | 0.19 | 4.90E+06 | 1.15 |
| 283Arg | 1.88 | 0.27 | 3.24E+06 | 1.72 |
| 286Glu | 42.40 | 4.12 | 3.55E+05 | 0.01 |

$k_{ex}$ = observed exchanged rate; $k_{ex}$ was obtain from the rate of decay. $R^2$ errors correspond deviation away from the regression line.
$k_{rc}$ = calculated exchange rate of that residue in a random coil, according to [14].
PF = Protection factors were calculated by dividing the exchange rate of the residue in a random coil by the observed hydrogen exchange rate ( $k_{rc}/k_{ex}$ ).

**Table S3.** Protection Factors for N235K p53.

| Residue | $k_{\text{ex}} (10^{-4})/\text{min}$ | Error | $k_{\text{rc}} (10^{-4})/\text{min}$ | PF ($10^6$) |
| --- | --- | --- | --- | --- |
| 110Arg | 1.10 | 0.09 | 2.24E+06 | 2.04 |
| 112Gly | 5.37 | 1.09 | 1.86E+06 | 0.35 |
| 132Lys | 5.29 | 1.24 | 3.09E+06 | 0.58 |
| 134Phe | 1.62 | 0.47 | 1.20E+06 | 0.74 |
| 135Cys | 5.20 | 1.25 | 7.76E+06 | 1.49 |
| 139Lys | 8.58 | 0.65 | 1.48E+06 | 0.17 |
| 141Cys | 0.67 | 0.58 | 1.07E+07 | 16.03 |
| 143Val | 0.21 | 0.05 | 1.86E+05 | 0.91 |
| 144Gln | 3.69 | 1.38 | 1.35E+06 | 0.37 |
| 145Leu | 1.65 | 0.28 | 6.76E+05 | 0.41 |
| 146Trp | 1.18 | 0.25 | 3.89E+05 | 0.33 |
| 156Arg | 0.33 | 0.04 | 3.09E+06 | 9.48 |
| 157Val | 0.17 | 0.03 | 5.37E+05 | 3.09 |
| 158Arg | 0.53 | 0.08 | 1.41E+06 | 2.68 |
| 159Ala | 0.20 | 0.03 | 2.69E+06 | 13.14 |
| 160Met | 0.26 | 0.05 | 1.59E+06 | 6.14 |
| 162Ile | 0.45 | 0.14 | 3.02E+05 | 0.67 |
| 163Tyr | 14.60 | 4.51 | 5.13E+05 | 0.04 |
| 195Ile | 0.69 | 0.09 | 1.86E+05 | 0.27 |
| 196Arg | 0.96 | 0.28 | 1.15E+06 | 1.20 |
| 204Glu | 1.54 | 0.10 | 3.63E+05 | 0.24 |
| 206Leu | 14.50 | 4.37 | 4.79E+05 | 0.03 |
| 215Ser | 4.58 | 0.53 | 5.25E+06 | 1.15 |
| 216Val | 0.21 | 0.03 | 6.46E+05 | 3.10 |
| 217Val | 0.29 | 0.05 | 2.34E+05 | 0.81 |
| 218Val | 0.14 | 0.02 | 2.34E+05 | 1.66 |
| 220Tyr | 1.43 | 0.10 | 5.01E+05 | 0.35 |
| 232Ile | 1.63 | 0.21 | 4.79E+05 | 0.29 |
| 234Tyr | 1.63 | 0.21 | 1.20E+06 | 0.74 |
| 235Lys | 1.53 | 0.16 | 2.19E+06 | 1.43 |
| 236Tyr | 1.17 | 0.21 | 1.15E+06 | 0.98 |
| 237Met | 0.04 | 0.10 | 1.78E+06 | 45.94 |
| 238Cys | 6.00 | 0.42 | 8.71E+06 | 1.45 |
| 252Leu | 0.52 | 0.08 | 2.51E+05 | 0.48 |
| 253Thr | 0.66 | 0.16 | 8.51E+05 | 1.29 |
| 254Ile | 0.68 | 0.10 | 4.79E+05 | 0.70 |
| 255Ile | 0.19 | 0.04 | 1.78E+05 | 0.94 |
| 256Thr | 0.09 | 0.03 | 8.13E+05 | 9.17 |
| 257Leu | 0.67 | 0.07 | 6.76E+05 | 1.01 |
| 258Glu | 0.23 | 0.03 | 3.09E+05 | 1.35 |
| 259Asp | 0.98 | 0.05 | 1.95E+07 | 19.94 |
| 265Leu | 0.35 | 0.09 | 2.63E+05 | 0.76 |
| 266Gly | 10.10 | 1.28 | 1.86E+06 | 0.18 |
| 268Asn | 0.25 | 0.03 | 8.32E+06 | 33.16 |
| 270Phe | 0.69 | 0.19 | 1.86E+06 | 2.70 |
| 272Val | 2.08 | 0.14 | 2.29E+05 | 0.11 |
| 273Arg | 1.09 | 0.22 | 1.41E+06 | 1.29 |
| 275Cys | 6.60 | 0.74 | 4.90E+06 | 0.74 |
| 283Arg | 1.85 | 0.18 | 3.24E+06 | 1.75 |

$k_{ex}$ = observed exchanged rate; $k_{ex}$ was obtain from the rate of decay. $R^2$ errors correspond deviation away from the regression line.
$k_{rc}$ = calculated exchange rate of that residue in a random coil, according to [14].
PF = Protection factors were calculated by dividing the exchange rate of the residue in a random coil by the observed hydrogen exchange rate ( $k_{rc}/k_{ex}$ ).

**Table S4.** Protection Factors for N239Y p53.

| Residue | $k_{\text{ex}} (10^{-4})/\text{min}$ | Error | $k_{\text{rc}} (10^{-4})/\text{min}$ | PF ($10^6$) |
| --- | --- | --- | --- | --- |
| 110Arg | 0.96 | 0.07 | 2.24E+06 | 2.33 |
| 112Gly | 4.41 | 0.34 | 1.86E+06 | 0.42 |
| 127Ser | 2.43 | 0.24 | 4.27E+06 | 1.76 |
| 132Lys | 1.93 | 0.09 | 3.09E+06 | 1.60 |
| 133Met | 1.67 | 0.27 | 2.09E+06 | 1.25 |
| 134Phe | 1.92 | 0.34 | 1.20E+06 | 0.63 |
| 135Cys | 1.36 | 0.06 | 7.76E+06 | 5.72 |
| 138Ala | 10.90 | 1.29 | 1.00E+06 | 0.09 |
| 141Cys | 2.94 | 0.29 | 1.07E+07 | 3.65 |
| 143Val | 0.53 | 0.06 | 1.86E+05 | 0.35 |
| 144Gln | 1.06 | 0.11 | 1.35E+06 | 1.27 |
| 145Leu | 0.66 | 0.08 | 6.76E+05 | 1.02 |
| 146Trp | 0.45 | 0.07 | 3.89E+05 | 0.87 |
| 156Arg | 0.65 | 0.08 | 3.09E+06 | 4.79 |
| 157Val | 0.35 | 0.05 | 5.37E+05 | 1.54 |
| 158Arg | 0.33 | 0.06 | 1.41E+06 | 4.25 |
| 159Ala | 0.26 | 0.04 | 2.69E+06 | 10.45 |
| 160Met | 0.41 | 0.05 | 1.59E+06 | 3.89 |
| 162Ile | 0.35 | 0.05 | 3.02E+05 | 0.85 |
| 163Tyr | 13.70 | 1.11 | 5.13E+05 | 0.04 |
| 193His | 2.46 | 0.33 | 2.57E+06 | 1.04 |
| 195Ile | 0.60 | 0.06 | 1.86E+05 | 0.31 |
| 196Arg | 1.02 | 0.12 | 1.15E+06 | 1.13 |
| 204Glu | 3.12 | 0.14 | 3.63E+05 | 0.12 |
| 206Leu | 0.80 | 0.14 | 4.79E+05 | 0.60 |
| 215Ser | 5.29 | 0.49 | 5.25E+06 | 0.99 |
| 216Val | 0.47 | 0.04 | 6.46E+05 | 1.38 |
| 217Val | 0.47 | 0.06 | 2.34E+05 | 0.49 |
| 218Val | 0.31 | 0.05 | 2.34E+05 | 0.76 |
| 220Tyr | 1.75 | 0.11 | 5.01E+05 | 0.29 |
| 230Thr | 46.57 | 6.14 | 4.90E+06 | 0.11 |
| 232Ile | 0.44 | 0.05 | 4.79E+05 | 1.08 |
| 234Tyr | 0.35 | 0.05 | 1.20E+06 | 3.46 |
| 235Asn | 0.81 | 0.07 | 5.62E+06 | 6.96 |
| 236Tyr | 0.47 | 0.06 | 1.82E+06 | 3.91 |
| 238Cys | 3.15 | 0.21 | 8.71E+06 | 2.77 |
| 252Leu | 0.44 | 0.06 | 2.51E+05 | 0.57 |
| 253Thr | 0.27 | 0.04 | 8.51E+05 | 3.19 |
| 254Ile | 0.69 | 0.11 | 4.79E+05 | 0.70 |
| 255Ile | 0.40 | 0.04 | 1.78E+05 | 0.45 |
| 256Thr | 0.30 | 0.04 | 8.13E+05 | 2.69 |
| 257Leu | 0.30 | 0.04 | 6.76E+05 | 2.24 |
| 258Glu | 0.35 | 0.05 | 3.09E+05 | 0.89 |
| 259Asp | 1.74 | 0.07 | 1.95E+07 | 11.18 |
| 265Leu | 0.60 | 0.13 | 2.63E+05 | 0.44 |
| 266Gly | 12.80 | 0.90 | 1.86E+06 | 0.15 |
| 268Asn | 0.35 | 0.06 | 8.32E+06 | 23.60 |
| 270Phe | 0.43 | 0.05 | 1.86E+06 | 4.32 |
| 272Val | 0.80 | 0.06 | 2.29E+05 | 0.29 |
| 273Arg | 0.98 | 0.11 | 1.41E+06 | 1.44 |
| 275Cys | 1.38 | 0.09 | 4.90E+06 | 3.54 |
| 283Arg | 3.26 | 0.08 | 3.24E+06 | 0.99 |
| 286Glu | 38.20 | 1.69 | 3.55E+05 | 0.01 |

$k_{ex}$ = observed exchanged rate; $k_{ex}$ was obtain from the rate of decay. $R^2$ errors correspond to deviations away from the regression line.
$k_{rc}$ = calculated exchange rate of that residue in a random coil, according to [14].
PF = Protection factors were calculated by dividing the exchange rate of the residue in a random coil by the observed hydrogen exchange rate ( $k_{rc}/k_{ex}$ ).

**Table S5.**
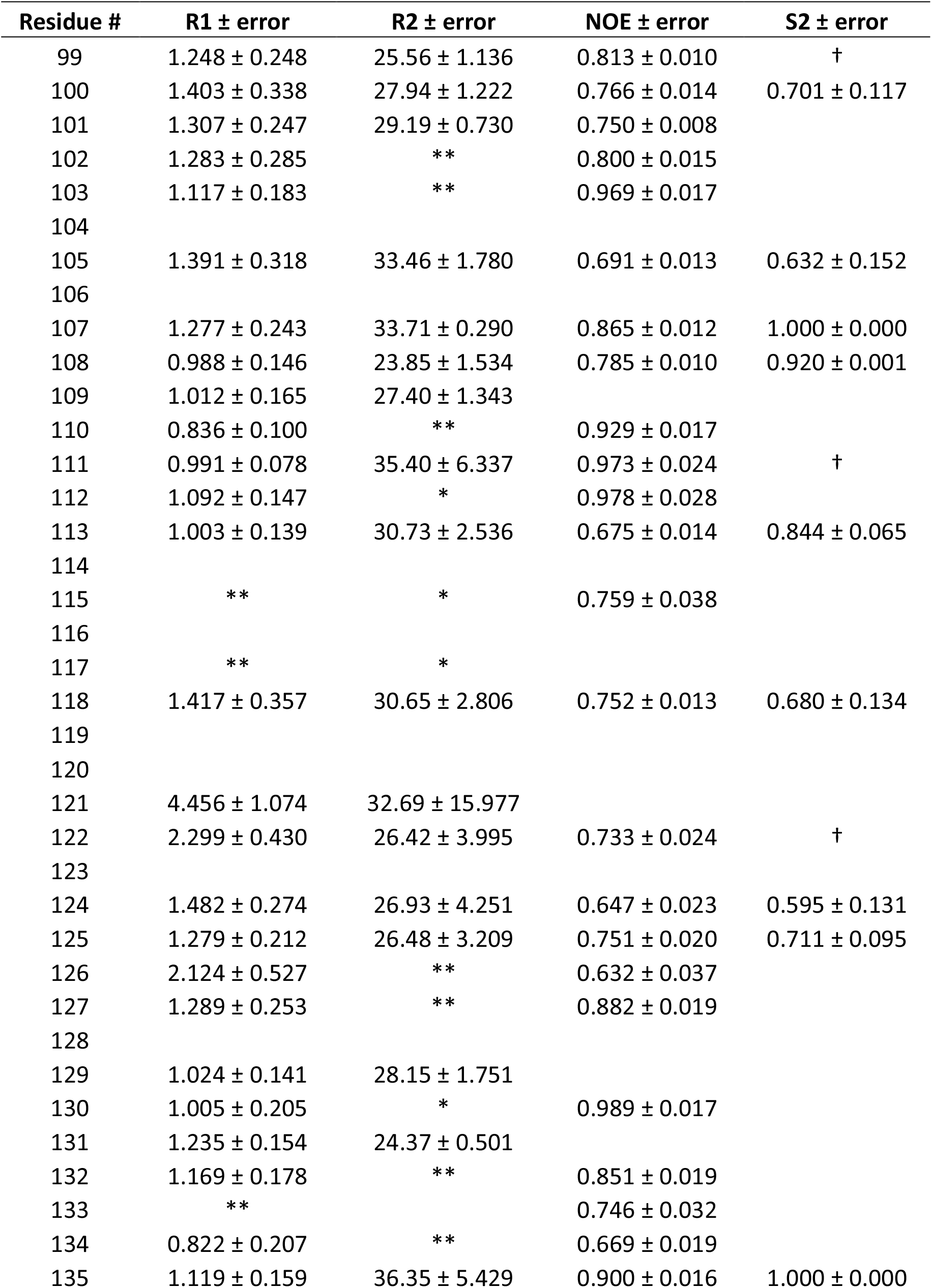
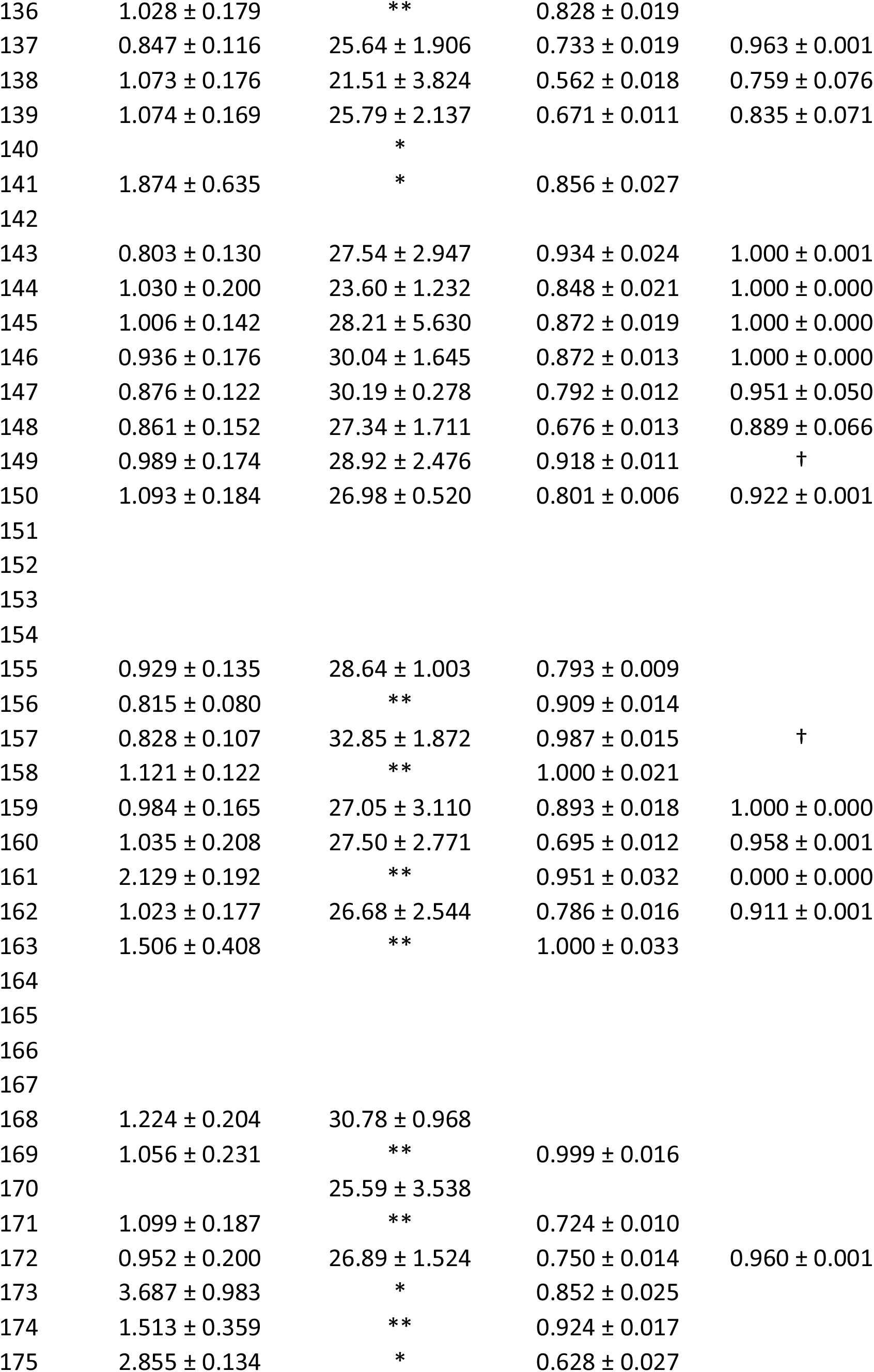
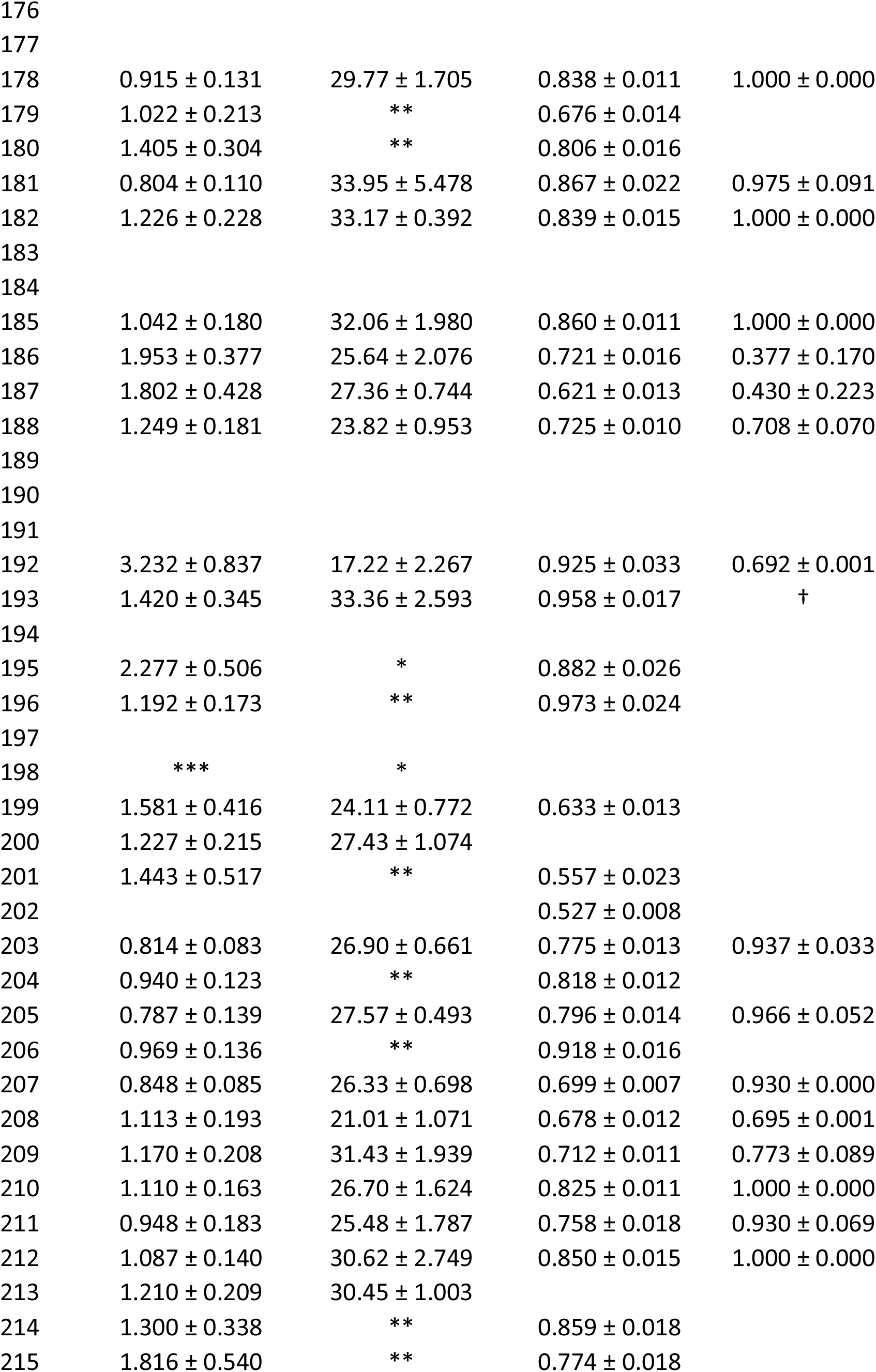
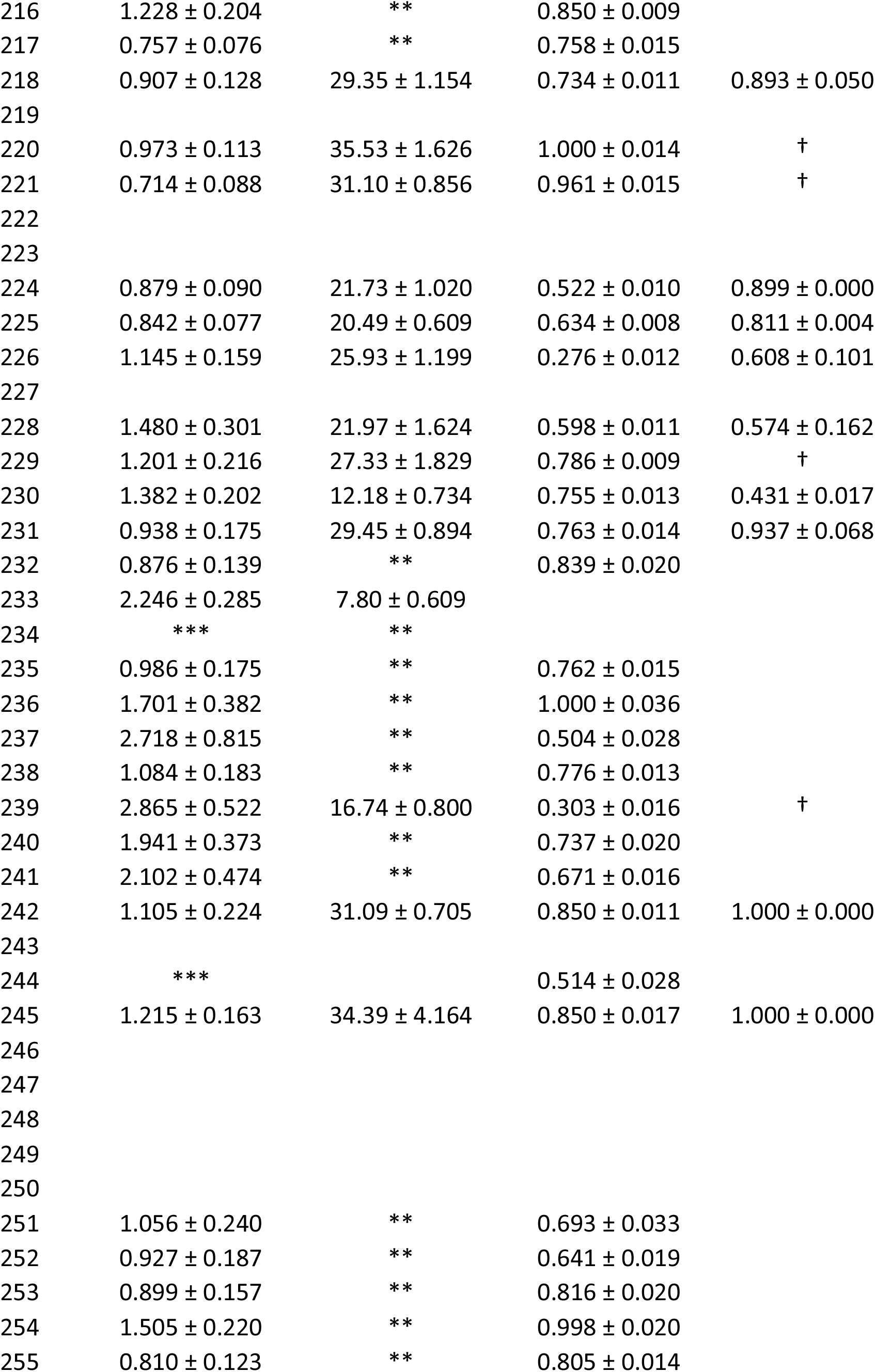
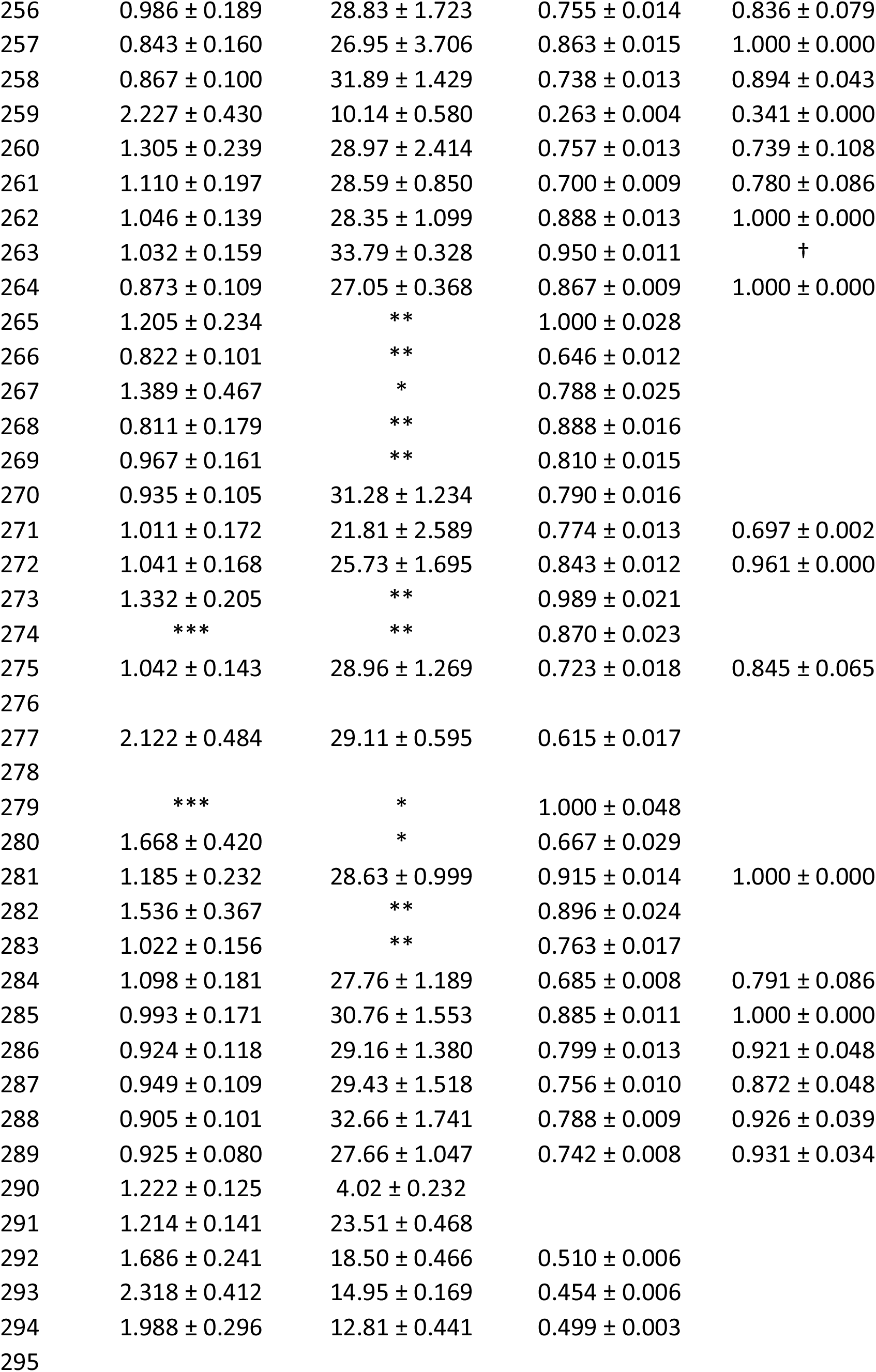
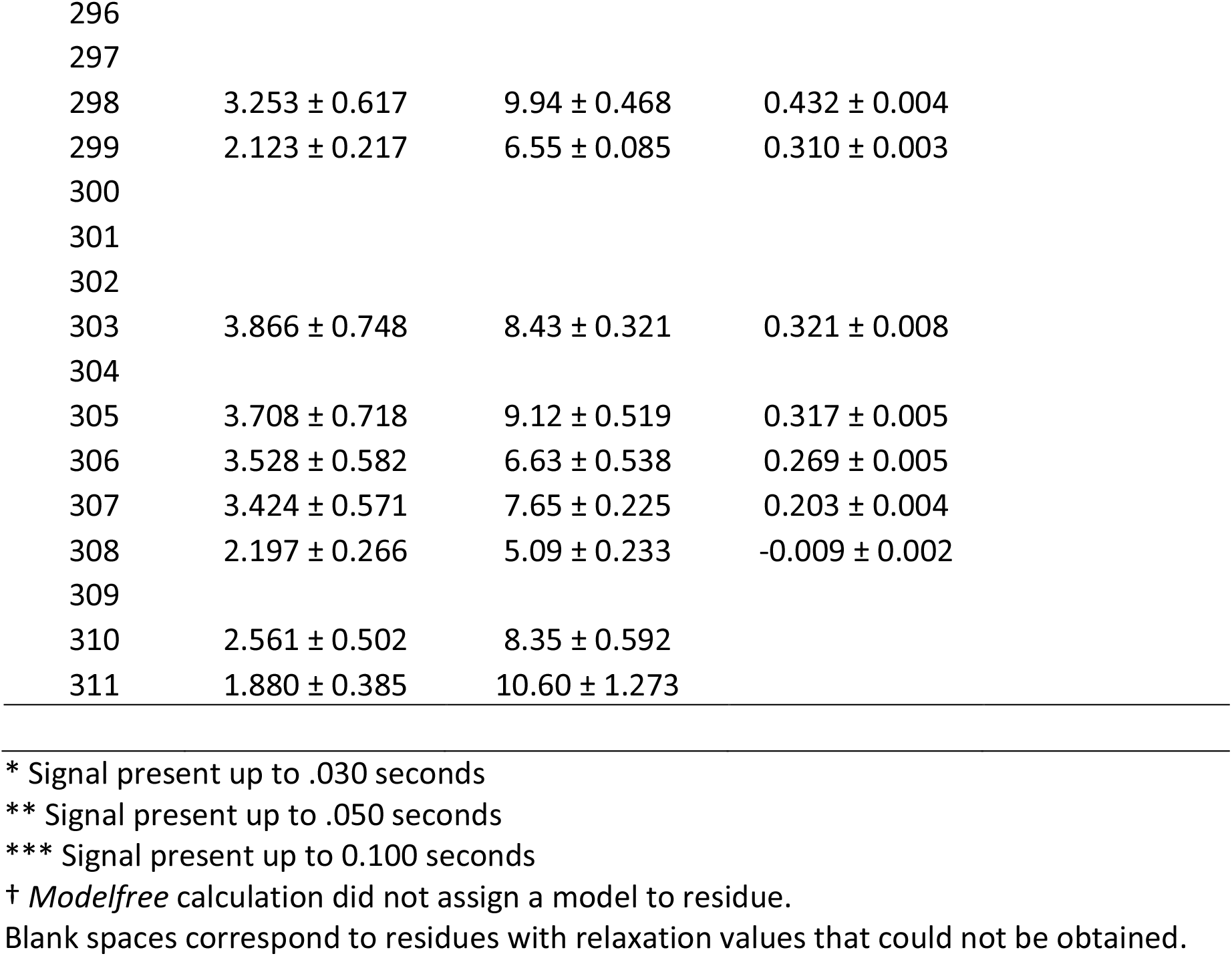
R1, R2, NOE, and S^2^ values for N235K p53 DBD at 800 MHz field strength.

| Residue # | R1 ± error | R2 ± error | NOE ± error | S2 ± error |
| --- | --- | --- | --- | --- |
| 99 | 1.248 ± 0.248 | 25.56 ± 1.136 | 0.813 ± 0.010 | † |
| 100 | 1.403 ± 0.338 | 27.94 ± 1.222 | 0.766 ± 0.014 | 0.701 ± 0.117 |
| 101 | 1.307 ± 0.247 | 29.19 ± 0.730 | 0.750 ± 0.008 |  |
| 102 | 1.283 ± 0.285 | ** | 0.800 ± 0.015 |  |
| 103 | 1.117 ± 0.183 | ** | 0.969 ± 0.017 |  |
| 104 |  |  |  |  |
| 105 | 1.391 ± 0.318 | 33.46 ± 1.780 | 0.691 ± 0.013 | 0.632 ± 0.152 |
| 106 |  |  |  |  |
| 107 | 1.277 ± 0.243 | 33.71 ± 0.290 | 0.865 ± 0.012 | 1.000 ± 0.000 |
| 108 | 0.988 ± 0.146 | 23.85 ± 1.534 | 0.785 ± 0.010 | 0.920 ± 0.001 |
| 109 | 1.012 ± 0.165 | 27.40 ± 1.343 |  |  |
| 110 | 0.836 ± 0.100 | ** | 0.929 ± 0.017 |  |
| 111 | 0.991 ± 0.078 | 35.40 ± 6.337 | 0.973 ± 0.024 | † |
| 112 | 1.092 ± 0.147 | * | 0.978 ± 0.028 |  |
| 113 | 1.003 ± 0.139 | 30.73 ± 2.536 | 0.675 ± 0.014 | 0.844 ± 0.065 |
| 114 |  |  |  |  |
| 115 | ** | * | 0.759 ± 0.038 |  |
| 116 |  |  |  |  |
| 117 | ** | * |  |  |
| 118 | 1.417 ± 0.357 | 30.65 ± 2.806 | 0.752 ± 0.013 | 0.680 ± 0.134 |
| 119 |  |  |  |  |
| 120 |  |  |  |  |
| 121 | 4.456 ± 1.074 | 32.69 ± 15.977 |  |  |
| 122 | 2.299 ± 0.430 | 26.42 ± 3.995 | 0.733 ± 0.024 | † |
| 123 |  |  |  |  |
| 124 | 1.482 ± 0.274 | 26.93 ± 4.251 | 0.647 ± 0.023 | 0.595 ± 0.131 |
| 125 | 1.279 ± 0.212 | 26.48 ± 3.209 | 0.751 ± 0.020 | 0.711 ± 0.095 |
| 126 | 2.124 ± 0.527 | ** | 0.632 ± 0.037 |  |
| 127 | 1.289 ± 0.253 | ** | 0.882 ± 0.019 |  |
| 128 |  |  |  |  |
| 129 | 1.024 ± 0.141 | 28.15 ± 1.751 |  |  |
| 130 | 1.005 ± 0.205 | * | 0.989 ± 0.017 |  |
| 131 | 1.235 ± 0.154 | 24.37 ± 0.501 |  |  |
| 132 | 1.169 ± 0.178 | ** | 0.851 ± 0.019 |  |
| 133 | ** |  | 0.746 ± 0.032 |  |
| 134 | 0.822 ± 0.207 | ** | 0.669 ± 0.019 |  |
| 135 | 1.119 ± 0.159 | 36.35 ± 5.429 | 0.900 ± 0.016 | 1.000 ± 0.000 |
| 136 | 1.028 ± 0.179 | ** | 0.828 ± 0.019 |  |
| 137 | 0.847 ± 0.116 | 25.64 ± 1.906 | 0.733 ± 0.019 | 0.963 ± 0.001 |
| 138 | 1.073 ± 0.176 | 21.51 ± 3.824 | 0.562 ± 0.018 | 0.759 ± 0.076 |
| 139 | 1.074 ± 0.169 | 25.79 ± 2.137 | 0.671 ± 0.011 | 0.835 ± 0.071 |
| 140 |  | * |  |  |
| 141 | 1.874 ± 0.635 | * | 0.856 ± 0.027 |  |
| 142 |  |  |  |  |
| 143 | 0.803 ± 0.130 | 27.54 ± 2.947 | 0.934 ± 0.024 | 1.000 ± 0.001 |
| 144 | 1.030 ± 0.200 | 23.60 ± 1.232 | 0.848 ± 0.021 | 1.000 ± 0.000 |
| 145 | 1.006 ± 0.142 | 28.21 ± 5.630 | 0.872 ± 0.019 | 1.000 ± 0.000 |
| 146 | 0.936 ± 0.176 | 30.04 ± 1.645 | 0.872 ± 0.013 | 1.000 ± 0.000 |
| 147 | 0.876 ± 0.122 | 30.19 ± 0.278 | 0.792 ± 0.012 | 0.951 ± 0.050 |
| 148 | 0.861 ± 0.152 | 27.34 ± 1.711 | 0.676 ± 0.013 | 0.889 ± 0.066 |
| 149 | 0.989 ± 0.174 | 28.92 ± 2.476 | 0.918 ± 0.011 | † |
| 150 | 1.093 ± 0.184 | 26.98 ± 0.520 | 0.801 ± 0.006 | 0.922 ± 0.001 |
| 151 |  |  |  |  |
| 152 |  |  |  |  |
| 153 |  |  |  |  |
| 154 |  |  |  |  |
| 155 | 0.929 ± 0.135 | 28.64 ± 1.003 | 0.793 ± 0.009 |  |
| 156 | 0.815 ± 0.080 | ** | 0.909 ± 0.014 |  |
| 157 | 0.828 ± 0.107 | 32.85 ± 1.872 | 0.987 ± 0.015 | † |
| 158 | 1.121 ± 0.122 | ** | 1.000 ± 0.021 |  |
| 159 | 0.984 ± 0.165 | 27.05 ± 3.110 | 0.893 ± 0.018 | 1.000 ± 0.000 |
| 160 | 1.035 ± 0.208 | 27.50 ± 2.771 | 0.695 ± 0.012 | 0.958 ± 0.001 |
| 161 | 2.129 ± 0.192 | ** | 0.951 ± 0.032 | 0.000 ± 0.000 |
| 162 | 1.023 ± 0.177 | 26.68 ± 2.544 | 0.786 ± 0.016 | 0.911 ± 0.001 |
| 163 | 1.506 ± 0.408 | ** | 1.000 ± 0.033 |  |
| 164 |  |  |  |  |
| 165 |  |  |  |  |
| 166 |  |  |  |  |
| 167 |  |  |  |  |
| 168 | 1.224 ± 0.204 | 30.78 ± 0.968 |  |  |
| 169 | 1.056 ± 0.231 | ** | 0.999 ± 0.016 |  |
| 170 |  | 25.59 ± 3.538 |  |  |
| 171 | 1.099 ± 0.187 | ** | 0.724 ± 0.010 |  |
| 172 | 0.952 ± 0.200 | 26.89 ± 1.524 | 0.750 ± 0.014 | 0.960 ± 0.001 |
| 173 | 3.687 ± 0.983 | * | 0.852 ± 0.025 |  |
| 174 | 1.513 ± 0.359 | ** | 0.924 ± 0.017 |  |
| 175 | 2.855 ± 0.134 | * | 0.628 ± 0.027 |  |
| 176 |  |  |  |  |
| 177 |  |  |  |  |
| 178 | $0.915 \pm 0.131$ | $29.77 \pm 1.705$ | $0.838 \pm 0.011$ | $1.000 \pm 0.000$ |
| 179 | $1.022 \pm 0.213$ | ** | $0.676 \pm 0.014$ | |
| 180 | $1.405 \pm 0.304$ | ** | $0.806 \pm 0.016$ | |
| 181 | $0.804 \pm 0.110$ | $33.95 \pm 5.478$ | $0.867 \pm 0.022$ | $0.975 \pm 0.091$ |
| 182 | $1.226 \pm 0.228$ | $33.17 \pm 0.392$ | $0.839 \pm 0.015$ | $1.000 \pm 0.000$ |
| 183 |  |  |  |  |
| 184 |  |  |  |  |
| 185 | $1.042 \pm 0.180$ | $32.06 \pm 1.980$ | $0.860 \pm 0.011$ | $1.000 \pm 0.000$ |
| 186 | $1.953 \pm 0.377$ | $25.64 \pm 2.076$ | $0.721 \pm 0.016$ | $0.377 \pm 0.170$ |
| 187 | $1.802 \pm 0.428$ | $27.36 \pm 0.744$ | $0.621 \pm 0.013$ | $0.430 \pm 0.223$ |
| 188 | $1.249 \pm 0.181$ | $23.82 \pm 0.953$ | $0.725 \pm 0.010$ | $0.708 \pm 0.070$ |
| 189 |  |  |  |  |
| 190 |  |  |  |  |
| 191 |  |  |  |  |
| 192 | $3.232 \pm 0.837$ | $17.22 \pm 2.267$ | $0.925 \pm 0.033$ | $0.692 \pm 0.001$ |
| 193 | $1.420 \pm 0.345$ | $33.36 \pm 2.593$ | $0.958 \pm 0.017$ | † |
| 194 |  |  |  |  |
| 195 | $2.277 \pm 0.506$ | * | $0.882 \pm 0.026$ | |
| 196 | $1.192 \pm 0.173$ | ** | $0.973 \pm 0.024$ | |
| 197 |  |  |  |  |
| 198 | *** | * |  |  |
| 199 | $1.581 \pm 0.416$ | $24.11 \pm 0.772$ | $0.633 \pm 0.013$ | |
| 200 | $1.227 \pm 0.215$ | $27.43 \pm 1.074$ | | |
| 201 | $1.443 \pm 0.517$ | ** | $0.557 \pm 0.023$ | |
| 202 | | | $0.527 \pm 0.008$ | |
| 203 | $0.814 \pm 0.083$ | $26.90 \pm 0.661$ | $0.775 \pm 0.013$ | $0.937 \pm 0.033$ |
| 204 | $0.940 \pm 0.123$ | ** | $0.818 \pm 0.012$ | |
| 205 | $0.787 \pm 0.139$ | $27.57 \pm 0.493$ | $0.796 \pm 0.014$ | $0.966 \pm 0.052$ |
| 206 | $0.969 \pm 0.136$ | ** | $0.918 \pm 0.016$ | |
| 207 | $0.848 \pm 0.085$ | $26.33 \pm 0.698$ | $0.699 \pm 0.007$ | $0.930 \pm 0.000$ |
| 208 | $1.113 \pm 0.193$ | $21.01 \pm 1.071$ | $0.678 \pm 0.012$ | $0.695 \pm 0.001$ |
| 209 | $1.170 \pm 0.208$ | $31.43 \pm 1.939$ | $0.712 \pm 0.011$ | $0.773 \pm 0.089$ |
| 210 | $1.110 \pm 0.163$ | $26.70 \pm 1.624$ | $0.825 \pm 0.011$ | $1.000 \pm 0.000$ |
| 211 | $0.948 \pm 0.183$ | $25.48 \pm 1.787$ | $0.758 \pm 0.018$ | $0.930 \pm 0.069$ |
| 212 | $1.087 \pm 0.140$ | $30.62 \pm 2.749$ | $0.850 \pm 0.015$ | $1.000 \pm 0.000$ |
| 213 | $1.210 \pm 0.209$ | $30.45 \pm 1.003$ | | |
| 214 | $1.300 \pm 0.338$ | ** | $0.859 \pm 0.018$ | |
| 215 | $1.816 \pm 0.540$ | ** | $0.774 \pm 0.018$ | |
| 216 | 1.228 ± 0.204 | ** | 0.850 ± 0.009 |  |
| 217 | 0.757 ± 0.076 | ** | 0.758 ± 0.015 |  |
| 218 | 0.907 ± 0.128 | 29.35 ± 1.154 | 0.734 ± 0.011 | 0.893 ± 0.050 |
| 219 |  |  |  |  |
| 220 | 0.973 ± 0.113 | 35.53 ± 1.626 | 1.000 ± 0.014 | † |
| 221 | 0.714 ± 0.088 | 31.10 ± 0.856 | 0.961 ± 0.015 | † |
| 222 |  |  |  |  |
| 223 |  |  |  |  |
| 224 | 0.879 ± 0.090 | 21.73 ± 1.020 | 0.522 ± 0.010 | 0.899 ± 0.000 |
| 225 | 0.842 ± 0.077 | 20.49 ± 0.609 | 0.634 ± 0.008 | 0.811 ± 0.004 |
| 226 | 1.145 ± 0.159 | 25.93 ± 1.199 | 0.276 ± 0.012 | 0.608 ± 0.101 |
| 227 |  |  |  |  |
| 228 | 1.480 ± 0.301 | 21.97 ± 1.624 | 0.598 ± 0.011 | 0.574 ± 0.162 |
| 229 | 1.201 ± 0.216 | 27.33 ± 1.829 | 0.786 ± 0.009 | † |
| 230 | 1.382 ± 0.202 | 12.18 ± 0.734 | 0.755 ± 0.013 | 0.431 ± 0.017 |
| 231 | 0.938 ± 0.175 | 29.45 ± 0.894 | 0.763 ± 0.014 | 0.937 ± 0.068 |
| 232 | 0.876 ± 0.139 | ** | 0.839 ± 0.020 |  |
| 233 | 2.246 ± 0.285 | 7.80 ± 0.609 |  |  |
| 234 | *** | ** |  |  |
| 235 | 0.986 ± 0.175 | ** | 0.762 ± 0.015 |  |
| 236 | 1.701 ± 0.382 | ** | 1.000 ± 0.036 |  |
| 237 | 2.718 ± 0.815 | ** | 0.504 ± 0.028 |  |
| 238 | 1.084 ± 0.183 | ** | 0.776 ± 0.013 |  |
| 239 | 2.865 ± 0.522 | 16.74 ± 0.800 | 0.303 ± 0.016 | † |
| 240 | 1.941 ± 0.373 | ** | 0.737 ± 0.020 |  |
| 241 | 2.102 ± 0.474 | ** | 0.671 ± 0.016 |  |
| 242 | 1.105 ± 0.224 | 31.09 ± 0.705 | 0.850 ± 0.011 | 1.000 ± 0.000 |
| 243 |  |  |  |  |
| 244 | *** |  | 0.514 ± 0.028 |  |
| 245 | 1.215 ± 0.163 | 34.39 ± 4.164 | 0.850 ± 0.017 | 1.000 ± 0.000 |
| 246 |  |  |  |  |
| 247 |  |  |  |  |
| 248 |  |  |  |  |
| 249 |  |  |  |  |
| 250 |  |  |  |  |
| 251 | 1.056 ± 0.240 | ** | 0.693 ± 0.033 |  |
| 252 | 0.927 ± 0.187 | ** | 0.641 ± 0.019 |  |
| 253 | 0.899 ± 0.157 | ** | 0.816 ± 0.020 |  |
| 254 | 1.505 ± 0.220 | ** | 0.998 ± 0.020 |  |
| 255 | 0.810 ± 0.123 | ** | 0.805 ± 0.014 |  |
| 256 | 0.986 ± 0.189 | 28.83 ± 1.723 | 0.755 ± 0.014 | 0.836 ± 0.079 |
| 257 | 0.843 ± 0.160 | 26.95 ± 3.706 | 0.863 ± 0.015 | 1.000 ± 0.000 |
| 258 | 0.867 ± 0.100 | 31.89 ± 1.429 | 0.738 ± 0.013 | 0.894 ± 0.043 |
| 259 | 2.227 ± 0.430 | 10.14 ± 0.580 | 0.263 ± 0.004 | 0.341 ± 0.000 |
| 260 | 1.305 ± 0.239 | 28.97 ± 2.414 | 0.757 ± 0.013 | 0.739 ± 0.108 |
| 261 | 1.110 ± 0.197 | 28.59 ± 0.850 | 0.700 ± 0.009 | 0.780 ± 0.086 |
| 262 | 1.046 ± 0.139 | 28.35 ± 1.099 | 0.888 ± 0.013 | 1.000 ± 0.000 |
| 263 | 1.032 ± 0.159 | 33.79 ± 0.328 | 0.950 ± 0.011 | † |
| 264 | 0.873 ± 0.109 | 27.05 ± 0.368 | 0.867 ± 0.009 | 1.000 ± 0.000 |
| 265 | 1.205 ± 0.234 | ** | 1.000 ± 0.028 |  |
| 266 | 0.822 ± 0.101 | ** | 0.646 ± 0.012 |  |
| 267 | 1.389 ± 0.467 | * | 0.788 ± 0.025 |  |
| 268 | 0.811 ± 0.179 | ** | 0.888 ± 0.016 |  |
| 269 | 0.967 ± 0.161 | ** | 0.810 ± 0.015 |  |
| 270 | 0.935 ± 0.105 | 31.28 ± 1.234 | 0.790 ± 0.016 |  |
| 271 | 1.011 ± 0.172 | 21.81 ± 2.589 | 0.774 ± 0.013 | 0.697 ± 0.002 |
| 272 | 1.041 ± 0.168 | 25.73 ± 1.695 | 0.843 ± 0.012 | 0.961 ± 0.000 |
| 273 | 1.332 ± 0.205 | ** | 0.989 ± 0.021 |  |
| 274 | *** | ** | 0.870 ± 0.023 |  |
| 275 | 1.042 ± 0.143 | 28.96 ± 1.269 | 0.723 ± 0.018 | 0.845 ± 0.065 |
| 276 |  |  |  |  |
| 277 | 2.122 ± 0.484 | 29.11 ± 0.595 | 0.615 ± 0.017 |  |
| 278 |  |  |  |  |
| 279 | *** | * | 1.000 ± 0.048 |  |
| 280 | 1.668 ± 0.420 | * | 0.667 ± 0.029 |  |
| 281 | 1.185 ± 0.232 | 28.63 ± 0.999 | 0.915 ± 0.014 | 1.000 ± 0.000 |
| 282 | 1.536 ± 0.367 | ** | 0.896 ± 0.024 |  |
| 283 | 1.022 ± 0.156 | ** | 0.763 ± 0.017 |  |
| 284 | 1.098 ± 0.181 | 27.76 ± 1.189 | 0.685 ± 0.008 | 0.791 ± 0.086 |
| 285 | 0.993 ± 0.171 | 30.76 ± 1.553 | 0.885 ± 0.011 | 1.000 ± 0.000 |
| 286 | 0.924 ± 0.118 | 29.16 ± 1.380 | 0.799 ± 0.013 | 0.921 ± 0.048 |
| 287 | 0.949 ± 0.109 | 29.43 ± 1.518 | 0.756 ± 0.010 | 0.872 ± 0.048 |
| 288 | 0.905 ± 0.101 | 32.66 ± 1.741 | 0.788 ± 0.009 | 0.926 ± 0.039 |
| 289 | 0.925 ± 0.080 | 27.66 ± 1.047 | 0.742 ± 0.008 | 0.931 ± 0.034 |
| 290 | 1.222 ± 0.125 | 4.02 ± 0.232 |  |  |
| 291 | 1.214 ± 0.141 | 23.51 ± 0.468 |  |  |
| 292 | 1.686 ± 0.241 | 18.50 ± 0.466 | 0.510 ± 0.006 |  |
| 293 | 2.318 ± 0.412 | 14.95 ± 0.169 | 0.454 ± 0.006 |  |
| 294 | 1.988 ± 0.296 | 12.81 ± 0.441 | 0.499 ± 0.003 |  |
| 295 |  |  |  |  |

|  |  |  |  |
| --- | --- | --- | --- |
| 296 |  |  |  |
| 297 |  |  |  |
| 298 | $3.253 \pm 0.617$ | $9.94 \pm 0.468$ | $0.432 \pm 0.004$ |
| 299 | $2.123 \pm 0.217$ | $6.55 \pm 0.085$ | $0.310 \pm 0.003$ |
| 300 |  |  |  |
| 301 |  |  |  |
| 302 |  |  |  |
| 303 | $3.866 \pm 0.748$ | $8.43 \pm 0.321$ | $0.321 \pm 0.008$ |
| 304 |  |  |  |
| 305 | $3.708 \pm 0.718$ | $9.12 \pm 0.519$ | $0.317 \pm 0.005$ |
| 306 | $3.528 \pm 0.582$ | $6.63 \pm 0.538$ | $0.269 \pm 0.005$ |
| 307 | $3.424 \pm 0.571$ | $7.65 \pm 0.225$ | $0.203 \pm 0.004$ |
| 308 | $2.197 \pm 0.266$ | $5.09 \pm 0.233$ | $-0.009 \pm 0.002$ |
| 309 |  |  |  |
| 310 | $2.561 \pm 0.502$ | $8.35 \pm 0.592$ | |
| 311 | $1.880 \pm 0.385$ | $10.60 \pm 1.273$ | |

\* Signal present up to .030 seconds
\*\* Signal present up to .050 seconds
\*\*\* Signal present up to 0.100 seconds
† *Modelfree* calculation did not assign a model to residue.
Blank spaces correspond to residues with relaxation values that could not be obtained.

**Table S6.**
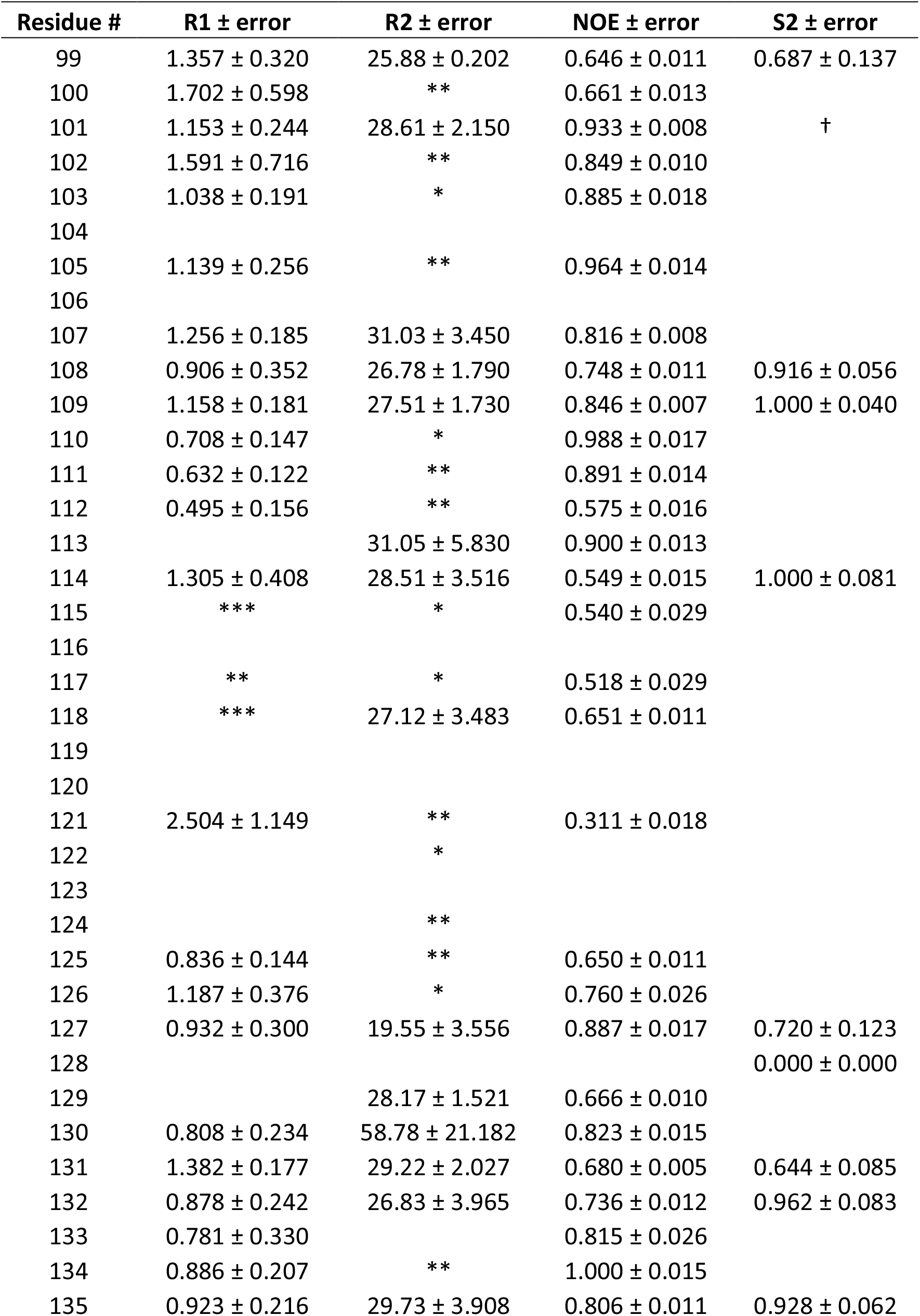
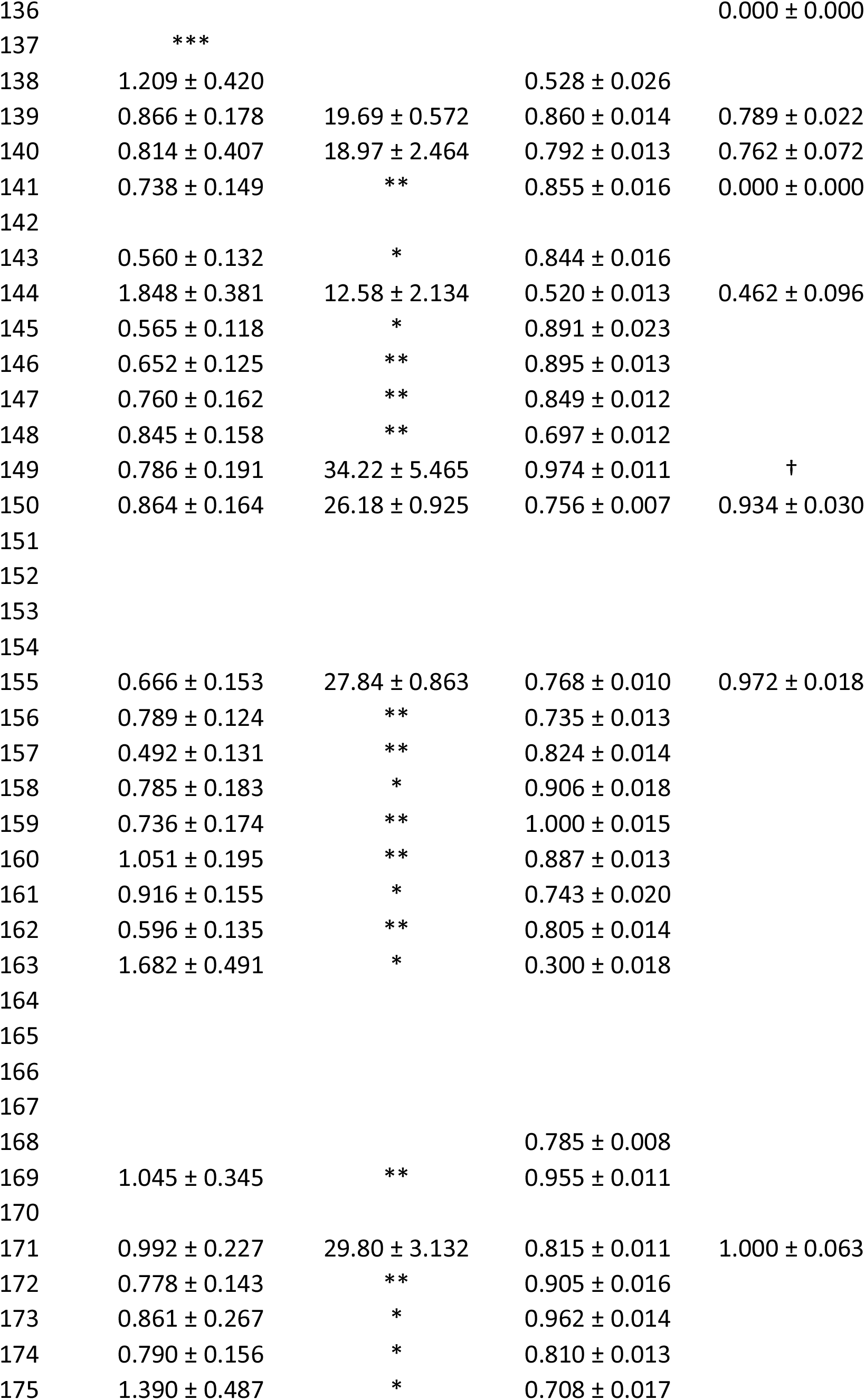
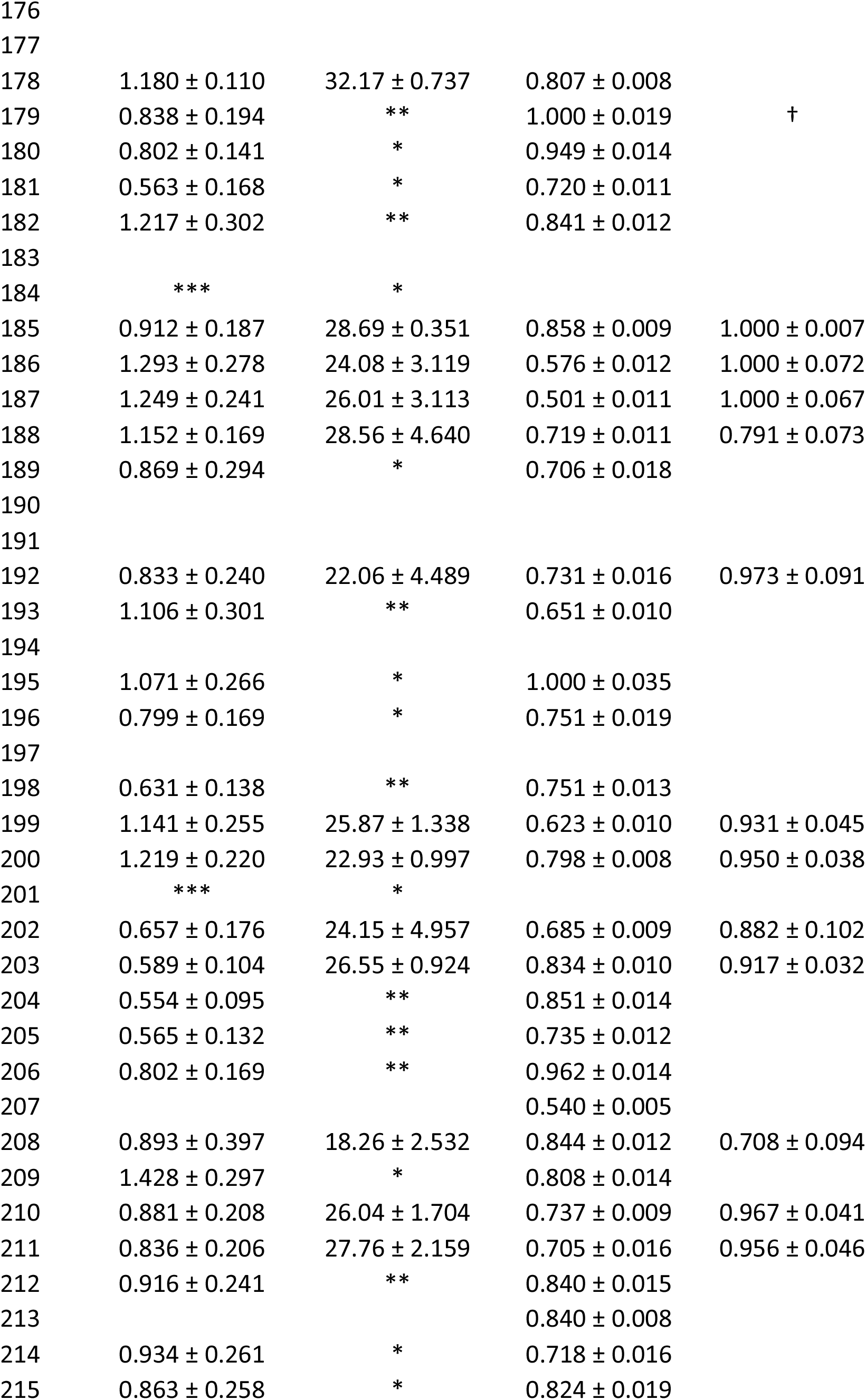
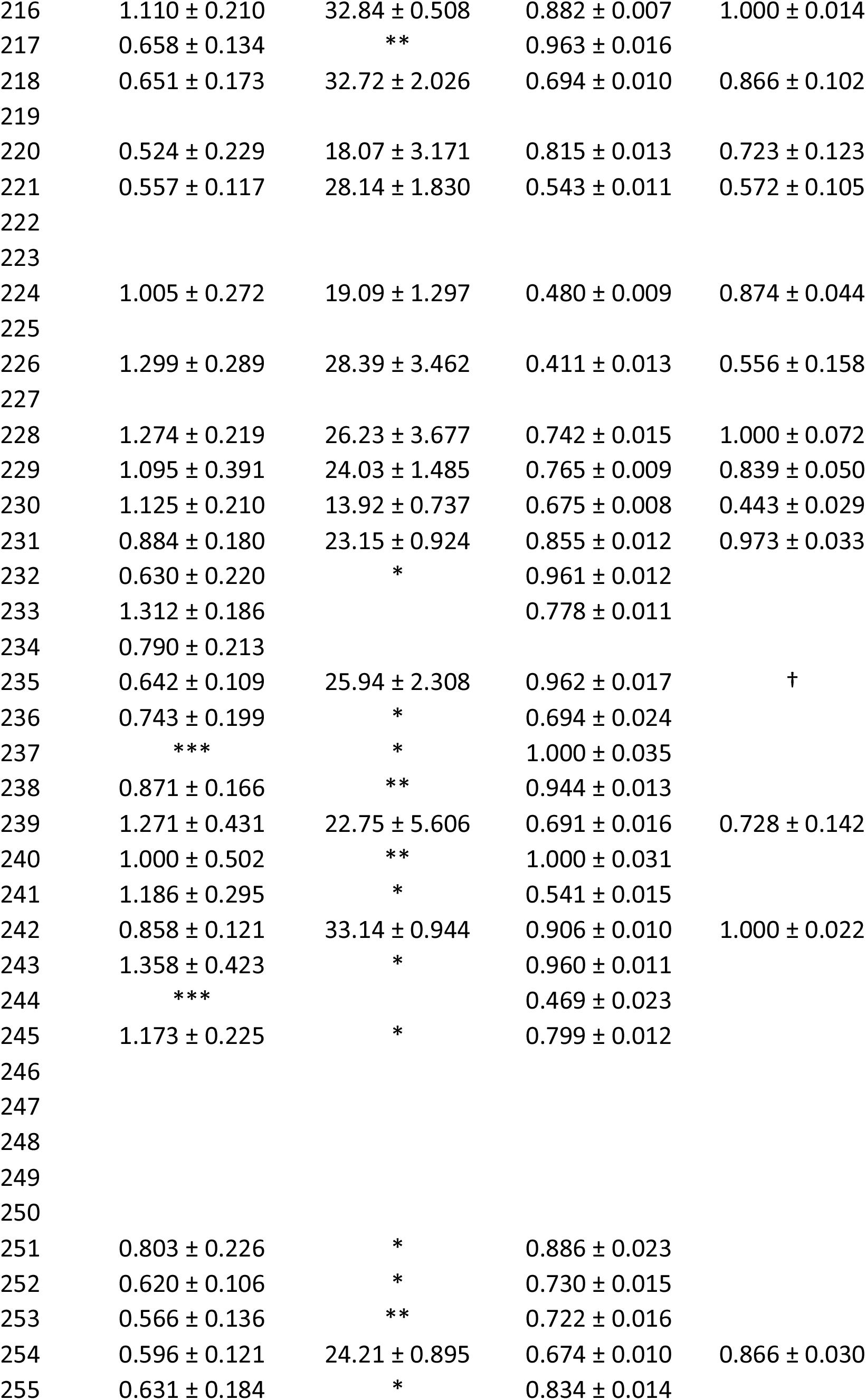
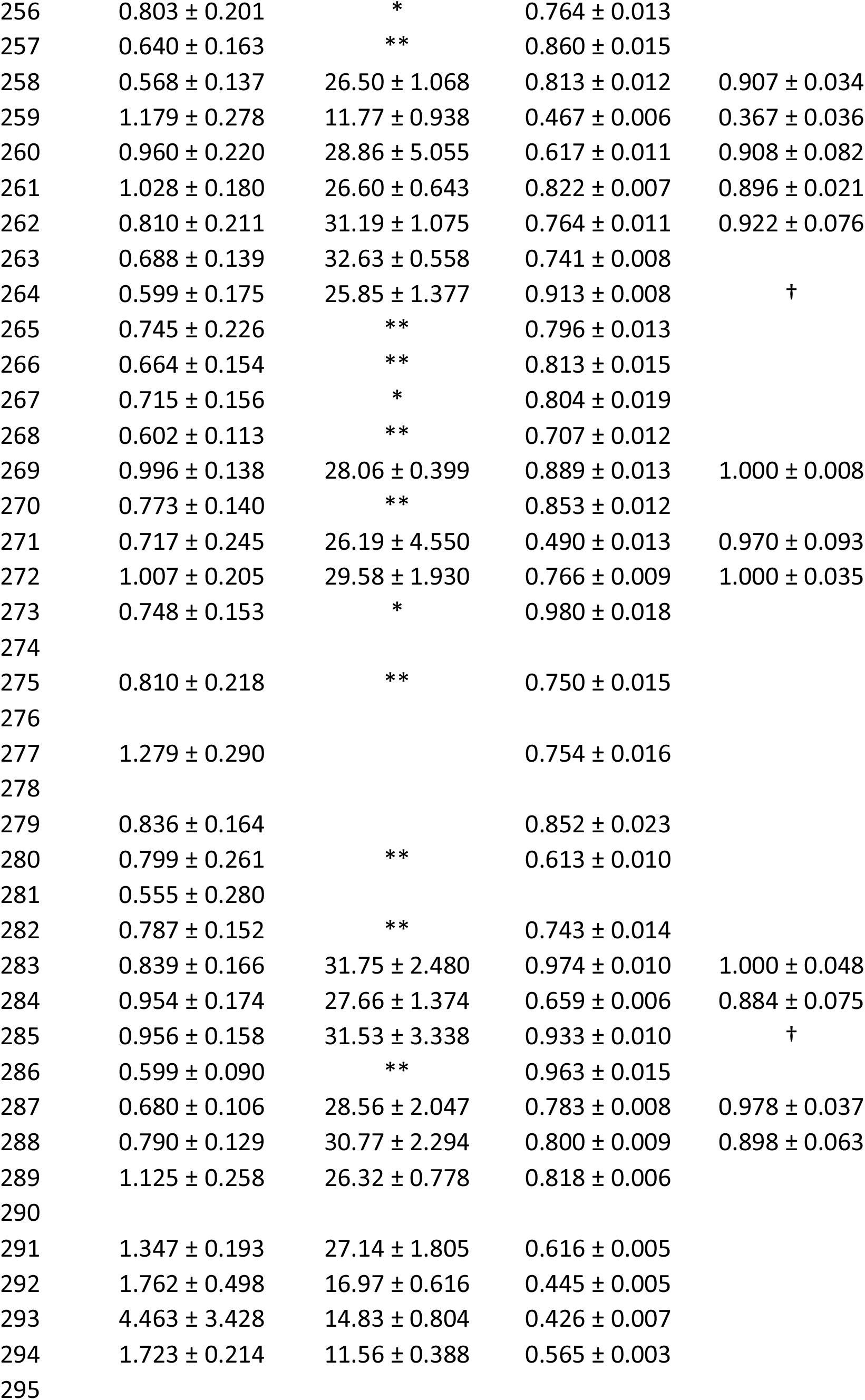
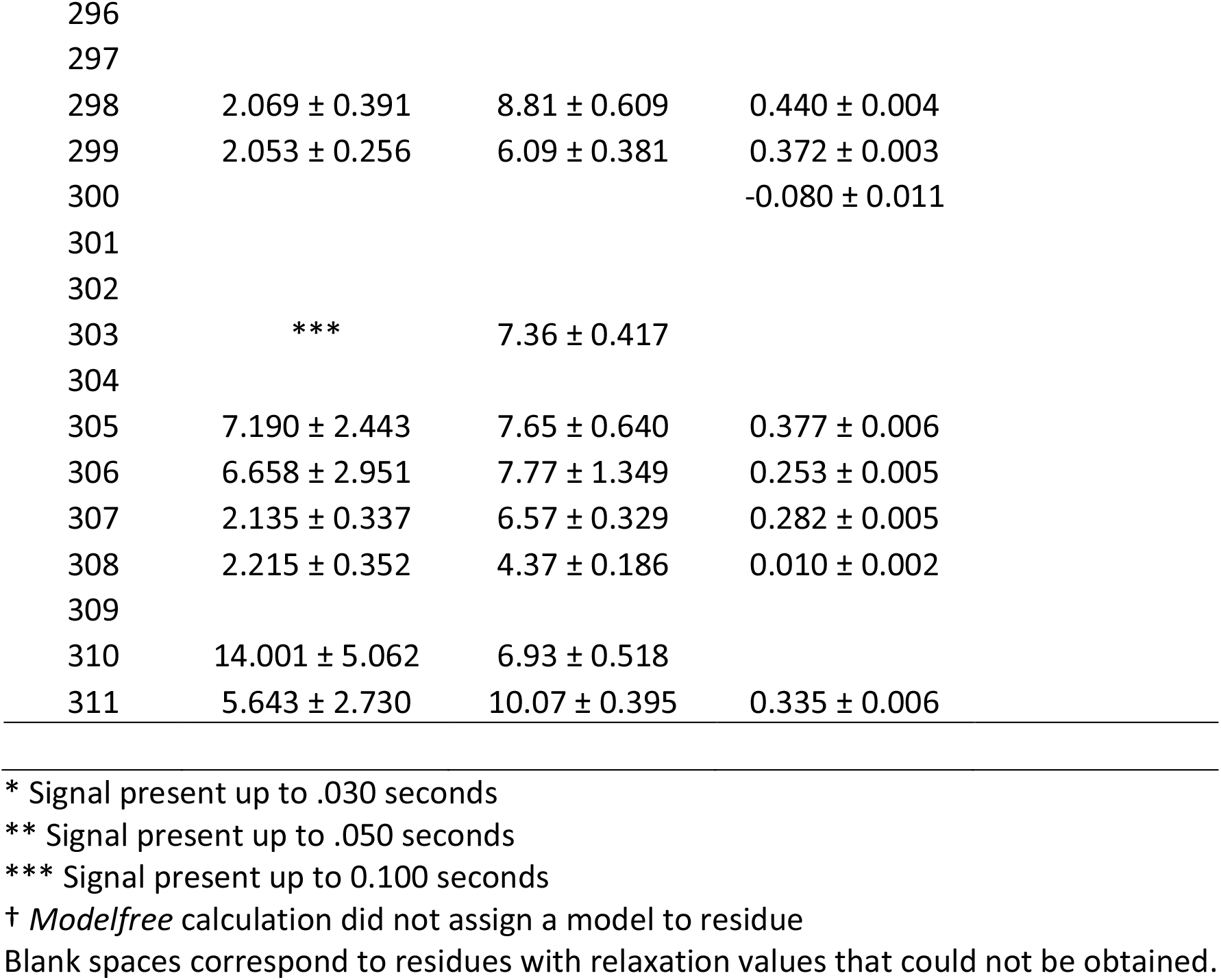
R1, R2, NOE, and S^2^ values for N239Y p53 DBD at 800 MHz field strength.

| Residue # | R1 ± error | R2 ± error | NOE ± error | S2 ± error |
| --- | --- | --- | --- | --- |
| 99 | 1.357 ± 0.320 | 25.88 ± 0.202 | 0.646 ± 0.011 | 0.687 ± 0.137 |
| 100 | 1.702 ± 0.598 | ** | 0.661 ± 0.013 |  |
| 101 | 1.153 ± 0.244 | 28.61 ± 2.150 | 0.933 ± 0.008 | † |
| 102 | 1.591 ± 0.716 | ** | 0.849 ± 0.010 |  |
| 103 | 1.038 ± 0.191 | * | 0.885 ± 0.018 |  |
| 104 |  |  |  |  |
| 105 | 1.139 ± 0.256 | ** | 0.964 ± 0.014 |  |
| 106 |  |  |  |  |
| 107 | 1.256 ± 0.185 | 31.03 ± 3.450 | 0.816 ± 0.008 |  |
| 108 | 0.906 ± 0.352 | 26.78 ± 1.790 | 0.748 ± 0.011 | 0.916 ± 0.056 |
| 109 | 1.158 ± 0.181 | 27.51 ± 1.730 | 0.846 ± 0.007 | 1.000 ± 0.040 |
| 110 | 0.708 ± 0.147 | * | 0.988 ± 0.017 |  |
| 111 | 0.632 ± 0.122 | ** | 0.891 ± 0.014 |  |
| 112 | 0.495 ± 0.156 | ** | 0.575 ± 0.016 |  |
| 113 |  | 31.05 ± 5.830 | 0.900 ± 0.013 |  |
| 114 | 1.305 ± 0.408 | 28.51 ± 3.516 | 0.549 ± 0.015 | 1.000 ± 0.081 |
| 115 | *** | * | 0.540 ± 0.029 |  |
| 116 |  |  |  |  |
| 117 | ** | * | 0.518 ± 0.029 |  |
| 118 | *** | 27.12 ± 3.483 | 0.651 ± 0.011 |  |
| 119 |  |  |  |  |
| 120 |  |  |  |  |
| 121 | 2.504 ± 1.149 | ** | 0.311 ± 0.018 |  |
| 122 |  | * |  |  |
| 123 |  |  |  |  |
| 124 |  | ** |  |  |
| 125 | 0.836 ± 0.144 | ** | 0.650 ± 0.011 |  |
| 126 | 1.187 ± 0.376 | * | 0.760 ± 0.026 |  |
| 127 | 0.932 ± 0.300 | 19.55 ± 3.556 | 0.887 ± 0.017 | 0.720 ± 0.123 |
| 128 |  |  |  | 0.000 ± 0.000 |
| 129 |  | 28.17 ± 1.521 | 0.666 ± 0.010 |  |
| 130 | 0.808 ± 0.234 | 58.78 ± 21.182 | 0.823 ± 0.015 |  |
| 131 | 1.382 ± 0.177 | 29.22 ± 2.027 | 0.680 ± 0.005 | 0.644 ± 0.085 |
| 132 | 0.878 ± 0.242 | 26.83 ± 3.965 | 0.736 ± 0.012 | 0.962 ± 0.083 |
| 133 | 0.781 ± 0.330 |  | 0.815 ± 0.026 |  |
| 134 | 0.886 ± 0.207 | ** | 1.000 ± 0.015 |  |
| 135 | 0.923 ± 0.216 | 29.73 ± 3.908 | 0.806 ± 0.011 | 0.928 ± 0.062 |
| 136 |  |  |  | 0.000 ± 0.000 |
| 137 | *** |  |  |  |
| 138 | 1.209 ± 0.420 |  | 0.528 ± 0.026 |  |
| 139 | 0.866 ± 0.178 | 19.69 ± 0.572 | 0.860 ± 0.014 | 0.789 ± 0.022 |
| 140 | 0.814 ± 0.407 | 18.97 ± 2.464 | 0.792 ± 0.013 | 0.762 ± 0.072 |
| 141 | 0.738 ± 0.149 | ** | 0.855 ± 0.016 | 0.000 ± 0.000 |
| 142 |  |  |  |  |
| 143 | 0.560 ± 0.132 | * | 0.844 ± 0.016 |  |
| 144 | 1.848 ± 0.381 | 12.58 ± 2.134 | 0.520 ± 0.013 | 0.462 ± 0.096 |
| 145 | 0.565 ± 0.118 | * | 0.891 ± 0.023 |  |
| 146 | 0.652 ± 0.125 | ** | 0.895 ± 0.013 |  |
| 147 | 0.760 ± 0.162 | ** | 0.849 ± 0.012 |  |
| 148 | 0.845 ± 0.158 | ** | 0.697 ± 0.012 |  |
| 149 | 0.786 ± 0.191 | 34.22 ± 5.465 | 0.974 ± 0.011 | † |
| 150 | 0.864 ± 0.164 | 26.18 ± 0.925 | 0.756 ± 0.007 | 0.934 ± 0.030 |
| 151 |  |  |  |  |
| 152 |  |  |  |  |
| 153 |  |  |  |  |
| 154 |  |  |  |  |
| 155 | 0.666 ± 0.153 | 27.84 ± 0.863 | 0.768 ± 0.010 | 0.972 ± 0.018 |
| 156 | 0.789 ± 0.124 | ** | 0.735 ± 0.013 |  |
| 157 | 0.492 ± 0.131 | ** | 0.824 ± 0.014 |  |
| 158 | 0.785 ± 0.183 | * | 0.906 ± 0.018 |  |
| 159 | 0.736 ± 0.174 | ** | 1.000 ± 0.015 |  |
| 160 | 1.051 ± 0.195 | ** | 0.887 ± 0.013 |  |
| 161 | 0.916 ± 0.155 | * | 0.743 ± 0.020 |  |
| 162 | 0.596 ± 0.135 | ** | 0.805 ± 0.014 |  |
| 163 | 1.682 ± 0.491 | * | 0.300 ± 0.018 |  |
| 164 |  |  |  |  |
| 165 |  |  |  |  |
| 166 |  |  |  |  |
| 167 |  |  |  |  |
| 168 |  |  | 0.785 ± 0.008 |  |
| 169 | 1.045 ± 0.345 | ** | 0.955 ± 0.011 |  |
| 170 |  |  |  |  |
| 171 | 0.992 ± 0.227 | 29.80 ± 3.132 | 0.815 ± 0.011 | 1.000 ± 0.063 |
| 172 | 0.778 ± 0.143 | ** | 0.905 ± 0.016 |  |
| 173 | 0.861 ± 0.267 | * | 0.962 ± 0.014 |  |
| 174 | 0.790 ± 0.156 | * | 0.810 ± 0.013 |  |
| 175 | 1.390 ± 0.487 | * | 0.708 ± 0.017 |  |
| 176 |  |  |  |  |
| 177 |  |  |  |  |
| 178 | $1.180 \pm 0.110$ | $32.17 \pm 0.737$ | $0.807 \pm 0.008$ | |
| 179 | $0.838 \pm 0.194$ | ** | $1.000 \pm 0.019$ | † |
| 180 | $0.802 \pm 0.141$ | * | $0.949 \pm 0.014$ | |
| 181 | $0.563 \pm 0.168$ | * | $0.720 \pm 0.011$ | |
| 182 | $1.217 \pm 0.302$ | ** | $0.841 \pm 0.012$ | |
| 183 |  |  |  |  |
| 184 | *** | * |  |  |
| 185 | $0.912 \pm 0.187$ | $28.69 \pm 0.351$ | $0.858 \pm 0.009$ | $1.000 \pm 0.007$ |
| 186 | $1.293 \pm 0.278$ | $24.08 \pm 3.119$ | $0.576 \pm 0.012$ | $1.000 \pm 0.072$ |
| 187 | $1.249 \pm 0.241$ | $26.01 \pm 3.113$ | $0.501 \pm 0.011$ | $1.000 \pm 0.067$ |
| 188 | $1.152 \pm 0.169$ | $28.56 \pm 4.640$ | $0.719 \pm 0.011$ | $0.791 \pm 0.073$ |
| 189 | $0.869 \pm 0.294$ | * | $0.706 \pm 0.018$ | |
| 190 |  |  |  |  |
| 191 |  |  |  |  |
| 192 | $0.833 \pm 0.240$ | $22.06 \pm 4.489$ | $0.731 \pm 0.016$ | $0.973 \pm 0.091$ |
| 193 | $1.106 \pm 0.301$ | ** | $0.651 \pm 0.010$ | |
| 194 |  |  |  |  |
| 195 | $1.071 \pm 0.266$ | * | $1.000 \pm 0.035$ | |
| 196 | $0.799 \pm 0.169$ | * | $0.751 \pm 0.019$ | |
| 197 |  |  |  |  |
| 198 | $0.631 \pm 0.138$ | ** | $0.751 \pm 0.013$ | |
| 199 | $1.141 \pm 0.255$ | $25.87 \pm 1.338$ | $0.623 \pm 0.010$ | $0.931 \pm 0.045$ |
| 200 | $1.219 \pm 0.220$ | $22.93 \pm 0.997$ | $0.798 \pm 0.008$ | $0.950 \pm 0.038$ |
| 201 | *** | * |  |  |
| 202 | $0.657 \pm 0.176$ | $24.15 \pm 4.957$ | $0.685 \pm 0.009$ | $0.882 \pm 0.102$ |
| 203 | $0.589 \pm 0.104$ | $26.55 \pm 0.924$ | $0.834 \pm 0.010$ | $0.917 \pm 0.032$ |
| 204 | $0.554 \pm 0.095$ | ** | $0.851 \pm 0.014$ | |
| 205 | $0.565 \pm 0.132$ | ** | $0.735 \pm 0.012$ | |
| 206 | $0.802 \pm 0.169$ | ** | $0.962 \pm 0.014$ | |
| 207 | | | $0.540 \pm 0.005$ | |
| 208 | $0.893 \pm 0.397$ | $18.26 \pm 2.532$ | $0.844 \pm 0.012$ | $0.708 \pm 0.094$ |
| 209 | $1.428 \pm 0.297$ | * | $0.808 \pm 0.014$ | |
| 210 | $0.881 \pm 0.208$ | $26.04 \pm 1.704$ | $0.737 \pm 0.009$ | $0.967 \pm 0.041$ |
| 211 | $0.836 \pm 0.206$ | $27.76 \pm 2.159$ | $0.705 \pm 0.016$ | $0.956 \pm 0.046$ |
| 212 | $0.916 \pm 0.241$ | ** | $0.840 \pm 0.015$ | |
| 213 | | | $0.840 \pm 0.008$ | |
| 214 | $0.934 \pm 0.261$ | * | $0.718 \pm 0.016$ | |
| 215 | $0.863 \pm 0.258$ | * | $0.824 \pm 0.019$ | |
| 216 | 1.110 ± 0.210 | 32.84 ± 0.508 | 0.882 ± 0.007 | 1.000 ± 0.014 |
| 217 | 0.658 ± 0.134 | ** | 0.963 ± 0.016 |  |
| 218 | 0.651 ± 0.173 | 32.72 ± 2.026 | 0.694 ± 0.010 | 0.866 ± 0.102 |
| 219 |  |  |  |  |
| 220 | 0.524 ± 0.229 | 18.07 ± 3.171 | 0.815 ± 0.013 | 0.723 ± 0.123 |
| 221 | 0.557 ± 0.117 | 28.14 ± 1.830 | 0.543 ± 0.011 | 0.572 ± 0.105 |
| 222 |  |  |  |  |
| 223 |  |  |  |  |
| 224 | 1.005 ± 0.272 | 19.09 ± 1.297 | 0.480 ± 0.009 | 0.874 ± 0.044 |
| 225 |  |  |  |  |
| 226 | 1.299 ± 0.289 | 28.39 ± 3.462 | 0.411 ± 0.013 | 0.556 ± 0.158 |
| 227 |  |  |  |  |
| 228 | 1.274 ± 0.219 | 26.23 ± 3.677 | 0.742 ± 0.015 | 1.000 ± 0.072 |
| 229 | 1.095 ± 0.391 | 24.03 ± 1.485 | 0.765 ± 0.009 | 0.839 ± 0.050 |
| 230 | 1.125 ± 0.210 | 13.92 ± 0.737 | 0.675 ± 0.008 | 0.443 ± 0.029 |
| 231 | 0.884 ± 0.180 | 23.15 ± 0.924 | 0.855 ± 0.012 | 0.973 ± 0.033 |
| 232 | 0.630 ± 0.220 | * | 0.961 ± 0.012 |  |
| 233 | 1.312 ± 0.186 |  | 0.778 ± 0.011 |  |
| 234 | 0.790 ± 0.213 |  |  |  |
| 235 | 0.642 ± 0.109 | 25.94 ± 2.308 | 0.962 ± 0.017 | † |
| 236 | 0.743 ± 0.199 | * | 0.694 ± 0.024 |  |
| 237 | *** | * | 1.000 ± 0.035 |  |
| 238 | 0.871 ± 0.166 | ** | 0.944 ± 0.013 |  |
| 239 | 1.271 ± 0.431 | 22.75 ± 5.606 | 0.691 ± 0.016 | 0.728 ± 0.142 |
| 240 | 1.000 ± 0.502 | ** | 1.000 ± 0.031 |  |
| 241 | 1.186 ± 0.295 | * | 0.541 ± 0.015 |  |
| 242 | 0.858 ± 0.121 | 33.14 ± 0.944 | 0.906 ± 0.010 | 1.000 ± 0.022 |
| 243 | 1.358 ± 0.423 | * | 0.960 ± 0.011 |  |
| 244 | *** |  | 0.469 ± 0.023 |  |
| 245 | 1.173 ± 0.225 | * | 0.799 ± 0.012 |  |
| 246 |  |  |  |  |
| 247 |  |  |  |  |
| 248 |  |  |  |  |
| 249 |  |  |  |  |
| 250 |  |  |  |  |
| 251 | 0.803 ± 0.226 | * | 0.886 ± 0.023 |  |
| 252 | 0.620 ± 0.106 | * | 0.730 ± 0.015 |  |
| 253 | 0.566 ± 0.136 | ** | 0.722 ± 0.016 |  |
| 254 | 0.596 ± 0.121 | 24.21 ± 0.895 | 0.674 ± 0.010 | 0.866 ± 0.030 |
| 255 | 0.631 ± 0.184 | * | 0.834 ± 0.014 |  |
| 256 | $0.803 \pm 0.201$ | * | $0.764 \pm 0.013$ | |
| 257 | $0.640 \pm 0.163$ | ** | $0.860 \pm 0.015$ | |
| 258 | $0.568 \pm 0.137$ | $26.50 \pm 1.068$ | $0.813 \pm 0.012$ | $0.907 \pm 0.034$ |
| 259 | $1.179 \pm 0.278$ | $11.77 \pm 0.938$ | $0.467 \pm 0.006$ | $0.367 \pm 0.036$ |
| 260 | $0.960 \pm 0.220$ | $28.86 \pm 5.055$ | $0.617 \pm 0.011$ | $0.908 \pm 0.082$ |
| 261 | $1.028 \pm 0.180$ | $26.60 \pm 0.643$ | $0.822 \pm 0.007$ | $0.896 \pm 0.021$ |
| 262 | $0.810 \pm 0.211$ | $31.19 \pm 1.075$ | $0.764 \pm 0.011$ | $0.922 \pm 0.076$ |
| 263 | $0.688 \pm 0.139$ | $32.63 \pm 0.558$ | $0.741 \pm 0.008$ | |
| 264 | $0.599 \pm 0.175$ | $25.85 \pm 1.377$ | $0.913 \pm 0.008$ | † |
| 265 | $0.745 \pm 0.226$ | ** | $0.796 \pm 0.013$ | |
| 266 | $0.664 \pm 0.154$ | ** | $0.813 \pm 0.015$ | |
| 267 | $0.715 \pm 0.156$ | * | $0.804 \pm 0.019$ | |
| 268 | $0.602 \pm 0.113$ | ** | $0.707 \pm 0.012$ | |
| 269 | $0.996 \pm 0.138$ | $28.06 \pm 0.399$ | $0.889 \pm 0.013$ | $1.000 \pm 0.008$ |
| 270 | $0.773 \pm 0.140$ | ** | $0.853 \pm 0.012$ | |
| 271 | $0.717 \pm 0.245$ | $26.19 \pm 4.550$ | $0.490 \pm 0.013$ | $0.970 \pm 0.093$ |
| 272 | $1.007 \pm 0.205$ | $29.58 \pm 1.930$ | $0.766 \pm 0.009$ | $1.000 \pm 0.035$ |
| 273 | $0.748 \pm 0.153$ | * | $0.980 \pm 0.018$ | |
| 274 |  |  |  |  |
| 275 | $0.810 \pm 0.218$ | ** | $0.750 \pm 0.015$ | |
| 276 |  |  |  |  |
| 277 | $1.279 \pm 0.290$ | | $0.754 \pm 0.016$ | |
| 278 |  |  |  |  |
| 279 | $0.836 \pm 0.164$ | | $0.852 \pm 0.023$ | |
| 280 | $0.799 \pm 0.261$ | ** | $0.613 \pm 0.010$ | |
| 281 | $0.555 \pm 0.280$ | | | |
| 282 | $0.787 \pm 0.152$ | ** | $0.743 \pm 0.014$ | |
| 283 | $0.839 \pm 0.166$ | $31.75 \pm 2.480$ | $0.974 \pm 0.010$ | $1.000 \pm 0.048$ |
| 284 | $0.954 \pm 0.174$ | $27.66 \pm 1.374$ | $0.659 \pm 0.006$ | $0.884 \pm 0.075$ |
| 285 | $0.956 \pm 0.158$ | $31.53 \pm 3.338$ | $0.933 \pm 0.010$ | † |
| 286 | $0.599 \pm 0.090$ | ** | $0.963 \pm 0.015$ | |
| 287 | $0.680 \pm 0.106$ | $28.56 \pm 2.047$ | $0.783 \pm 0.008$ | $0.978 \pm 0.037$ |
| 288 | $0.790 \pm 0.129$ | $30.77 \pm 2.294$ | $0.800 \pm 0.009$ | $0.898 \pm 0.063$ |
| 289 | $1.125 \pm 0.258$ | $26.32 \pm 0.778$ | $0.818 \pm 0.006$ | |
| 290 |  |  |  |  |
| 291 | $1.347 \pm 0.193$ | $27.14 \pm 1.805$ | $0.616 \pm 0.005$ | |
| 292 | $1.762 \pm 0.498$ | $16.97 \pm 0.616$ | $0.445 \pm 0.005$ | |
| 293 | $4.463 \pm 3.428$ | $14.83 \pm 0.804$ | $0.426 \pm 0.007$ | |
| 294 | $1.723 \pm 0.214$ | $11.56 \pm 0.388$ | $0.565 \pm 0.003$ | |
| 295 |  |  |  |  |

|  |  |  |  |
| --- | --- | --- | --- |
| 296 |  |  |  |
| 297 |  |  |  |
| 298 | 2.069 ± 0.391 | 8.81 ± 0.609 | 0.440 ± 0.004 |
| 299 | 2.053 ± 0.256 | 6.09 ± 0.381 | 0.372 ± 0.003 |
| 300 |  |  | -0.080 ± 0.011 |
| 301 |  |  |  |
| 302 |  |  |  |
| 303 | *** | 7.36 ± 0.417 |  |
| 304 |  |  |  |
| 305 | 7.190 ± 2.443 | 7.65 ± 0.640 | 0.377 ± 0.006 |
| 306 | 6.658 ± 2.951 | 7.77 ± 1.349 | 0.253 ± 0.005 |
| 307 | 2.135 ± 0.337 | 6.57 ± 0.329 | 0.282 ± 0.005 |
| 308 | 2.215 ± 0.352 | 4.37 ± 0.186 | 0.010 ± 0.002 |
| 309 |  |  |  |
| 310 | 14.001 ± 5.062 | 6.93 ± 0.518 |  |
| 311 | 5.643 ± 2.730 | 10.07 ± 0.395 | 0.335 ± 0.006 |

\* Signal present up to .030 seconds
\*\* Signal present up to .050 seconds
\*\*\* Signal present up to 0.100 seconds
† *Modelfree* calculation did not assign a model to residue
Blank spaces correspond to residues with relaxation values that could not be obtained.

